# Ultrasensitive voltage imaging reveals distinct electrical microdomains in neurons

**DOI:** 10.64898/2026.05.27.728040

**Authors:** Yukun Alex Hao, Lorna Louise Jayne, Sungmoo Lee, Mia Nicole Dittrich, Manze Zhang, Simon Haziza, Imane Bendifallah, Ruth R. Sims, Boris Bouazza-Arostegui, Alexander D. White, John Kochalka, Yu Wang, Maedeh Seyedolmohadesin, Adrian Negrean, Zhaoyang Li, Collin Chiu, Kaspar Podgorski, Jun B. Ding, Karl Deisseroth, Rafael Yuste, Valentina Emiliani, Mark J. Schnitzer, Michael Z. Lin, Thomas R. Clandinin

**Affiliations:** Department of Neurobiology, Stanford University, Stanford, CA, USA; Department of Bioengineering, Stanford University, Stanford, CA, USA; Department of Biology, Stanford University, Stanford, CA, USA; Center for Cracking the Neural Code, Stanford University, Stanford, CA, USA; Institut de la Vision, Sorbonne Université, INSERM, CNRS, Paris, France; Department of Biological Sciences, Columbia University, New York, NY, USA; Neurotechnology Center, Columbia University, New York, NY, USA; Allen Institute for Neural Dynamics, Seattle, WA, USA; Department of Biochemistry, Stanford University, Stanford, CA, USA; Department of Neurosurgery, Stanford University, Stanford, CA, USA; Department of Neurology and Neurological Sciences, Stanford University, Stanford, CA, USA; Department of Psychiatry and Behavioral Sciences, Stanford University, Stanford, CA, USA; Howard Hughes Medical Institute, Stanford University, Stanford, CA, USA; Department of Applied Physics, Stanford University, Stanford, CA, USA; Department of Chemical and Systems Biology, Stanford University, Stanford, CA, USA; Chan-Zuckerberg Biohub, San Francisco, CA

## Abstract

For the brain to compute, electrical signals must propagate over the membranes of individual neurons, connecting synaptic inputs to synaptic outputs^1^. Complex neuronal morphologies coupled with the spatial organization of synaptic inputs and outputs enable diverse voltage transformations that underlie cell-type specific computations^2,3^. However, measuring these transformations *in vivo* has remained challenging, leaving a crucial gap in our mechanistic understanding of single neuron computation. Here, we develop ASAP7y, a genetically encoded voltage indicator with unprecedented subthreshold sensitivity and expanded excitation compatibility in both mice and flies. We leveraged ASAP7y combined with two-photon random-access microscopy to record sensory stimulus-evoked voltage dynamics with millisecond, subcellular, and subthreshold resolution along the neurites of individual neurons in *Drosophila*. We found remarkable heterogeneity in voltage propagation across cell-types, delineating a fundamental axis of electrical diversity. Leveraging a nanoscale EM reconstruction of the visual system^4^, we modeled the electrotonic properties of single neurons spanning 717 cell types, revealing how morphology shapes voltage transformations. Finally, we demonstrate that confined voltage propagation creates substrates for local computation, producing subcellular domains with distinct feature selectivity across multiple cell types. These results provide mechanistic insight into how critical single neuron computations arise and reveal parallel processing in single neurons.

## Introduction

Neuronal computation relies on both chemical and electrical signals. Neurotransmitters, the chemical inputs to neurons, alter the permeability of neuronal membranes to ions, generating local voltage changes that propagate across each cell’s surface to shape synaptic output. Complex neuronal morphologies, the spatial distribution of input and output synapses, as well as the specific patterns of ion channel expression enable diverse voltage transformations that inform cell-type and input specific computations. As Ramon y Cajal first described, neurons exist in a diverse array of morphological types^5,6^. Subsequent molecular and electrophysiological studies have revealed remarkable complexity in the distributions of ion channels and the locations of synaptic inputs and outputs^2,3,7^. Thus, understanding how changes in membrane potential propagate from input to output synapses across cell types is central to a mechanistic understanding of single-neuron computation.

Both theory and experiment have demonstrated that active and passive neuronal properties enable diverse transformations of membrane potential. For example, the Hodgkin-Huxley model of the action potential quantitatively describes the biophysical substrates of active regeneration and propagation, while cable theory describes how neuronal morphology filters electrical signals as they passively propagate^8-10^. Although many such transformations of membrane potential have been mapped *in vitro^11^*, it has been challenging to directly measure how changes in membrane potential emerge and propagate between input and output synapses within single neurons *in vivo.* Thus, how complex morphologies, ion channel properties, and the spatial distribution of synapses construct cell-type specific single-neuron computations *in vivo* remains incompletely understood.

The challenge of measuring voltage dynamics *in vivo* arises from the intersection of three distinct features of neuronal processing: (1) neurites are often thin, limiting the use of intracellular electrodes; (2) voltage propagation over the surface of individual neurons is rapid, necessitating high temporal resolution; and (3) the small amplitude of subthreshold voltage signals requires a recording method with high sensitivity. Imaging-based methods to record changes in neuronal membrane potential have the potential to address these challenges, particularly through the development of fast imaging modalities, including compatibility with two-photon excitation, and through the development of new genetically encoded voltage indicators (GEVIs). Recent work combining two-photon imaging and GEVIs has been applied to record neuronal voltage *in vivo^12,13^*. These studies revealed spine-specific voltage events, input-output relationships in neuronal populations, and cell-type and compartment-specific transformations of voltage signals into intracellular calcium^14-17^. However, these recordings have been limited by signal-to-noise ratio (SNR), largely preventing sustained recordings of subthreshold, subcellular voltage dynamics on single trials. Thus, achieving reliable single-trial sensitivity in individual neurons in response to physiologically relevant stimuli over long recordings *in vivo* remains a critical goal for dissecting single neuron computations.

The visual system of *Drosophila melanogaster* provides a powerful model for investigating the voltage dynamics underlying information processing in single neurons^18^. This system is highly structured, maintaining a columnar, retinotopic arrangement across its major ganglia: the lamina, medulla, lobula, and lobula plate. These regions process visual information in a hierarchical manner, using broadly conserved algorithms to transform photoreceptor inputs into specialized signals that guide behavior. These ganglia include many morphologically and molecularly diverse neuron types^19,20^, and previous work has revealed the importance of local and single-neuron computations in the emergence of feature selectivity^14,17,21-35^. Moreover, given that the morphologies of each cell type are highly stereotyped within and across animals^36-39^, and are often entirely optically accessible, this system allows feature selective responses to be mapped across subcellular compartments and cell types *in vivo*. Finally, the genetic tractability of this system enables selective, sparse access to genetically defined cell-types^40^, while the recently published connectomes^4,41,42^ offer nanoscale reconstructions of the morphologies and subcellular synaptic connections of single neurons, facilitating realistic biophysical modeling.

Here we develop a highly sensitive GEVI, ASAP7y, that reports bidirectional changes in membrane potential and use it to capture single-trial multisite voltage responses with millivolt precision and kilohertz resolution across a diversity of cell types *in vivo*. By integrating these measurements with connectome-informed biophysical modeling, we reveal morphological substrates that underpin remarkable heterogeneity in voltage propagation across cell types. These studies reveal a continuum of cell-type specific tuning of voltage propagation that greatly expands the computational capacity of single neurons.

## Results

### Development of a new GEVI with higher subthreshold responsivity and broader two-photon compatibility

We conducted several rounds of structure-based single-site and combinatorial mutagenesis to obtain ASAP7y. When excited in the two-photon regime, ASAP7y showed peak brightness at 1000 nm, while maintaining >50% max brightness between 932 nm and 1038 nm (Fig. 1a). We compared ASAP7y to its predecessor, ASAP5, in response amplitude and kinetics by voltage-clamp electrophysiology. When normalized to fluorescence at resting membrane potential, ASAP7y displayed a relative fluorescence change (Δ*F*/*F*_0_) of –80% across the physiological voltage range from –70 mV to 30 mV, and a voltage-fluorescence conversion ratio of 2.7%/mV around the resting membrane potential, a two-fold increase in responsivity compared to ASAP5 (Fig. 1b). Across all voltage steps, ASAP7y demonstrated a full dynamic range of 18 compared to 3.5 for ASAP5 (Extended Data Fig. 1). Onset kinetics of ASAP7y exhibited a time constant of 3.0 ms at 25°C, and 0.9 ms at 37°C (Extended Data Fig. 1). ASAP7y responded to a 3-mV excitatory postsynaptic potential (EPSP) waveform with –6% Δ*F*/*F*_0_, double the response of ASAP5, and to a 100-mV action potential (AP) waveform with 62% ΔF/F_0_, a ∼50% increase over ASAP5 (Extended Data Fig. 1).

**Figure 1.**
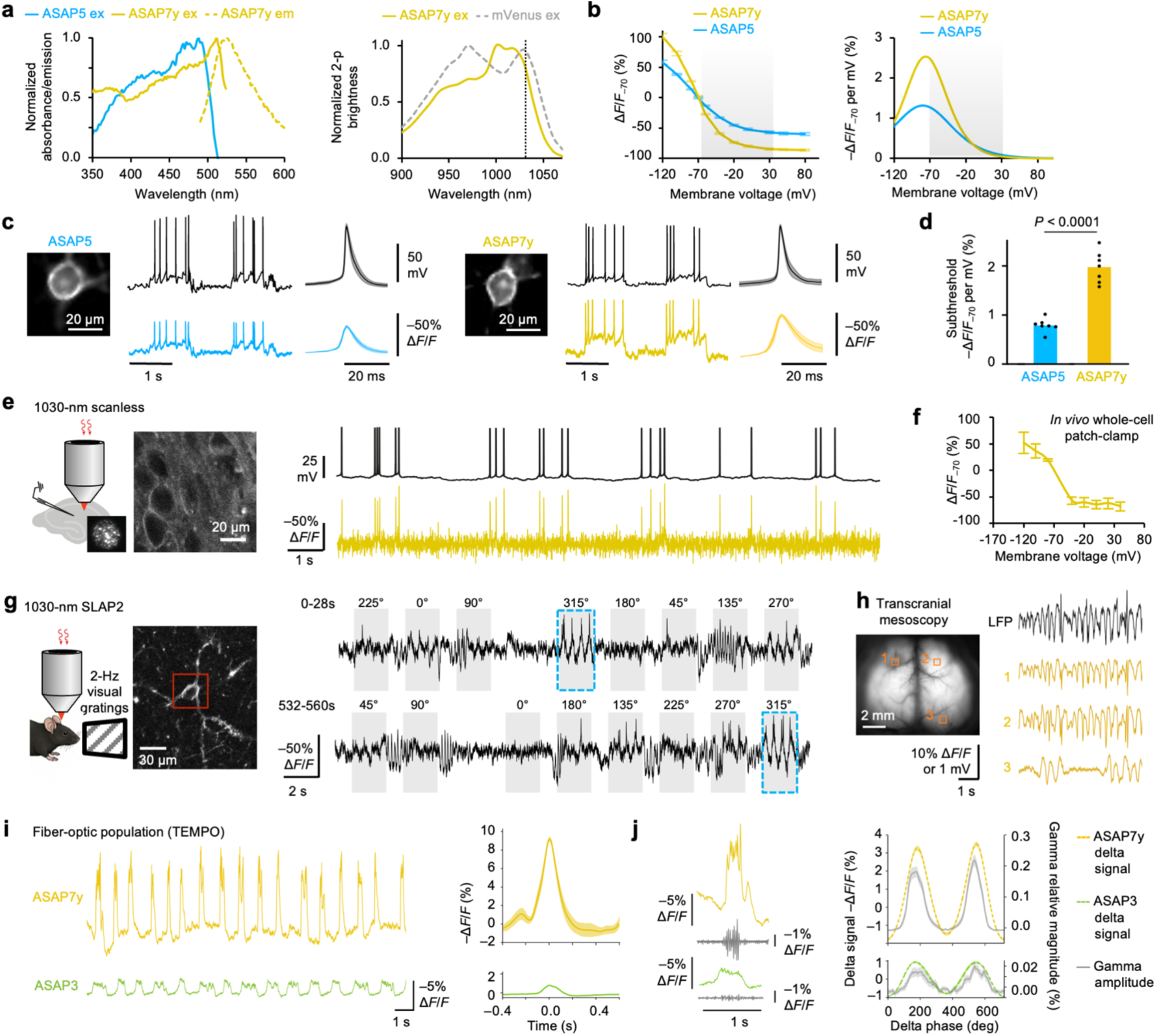
An improved red-shifted voltage indicator, ASAP7y, can be widely applied across imaging modalities. **a**, Left, one-photon excitation spectra of ASAP5 and ASAP7y, and emission spectrum of ASAP7y. Right, normalized 2-photon brightness of ASAP7y. A protein with mVenus YFP in place of cpGFP in ASAP7y was used for reference. Dashed line, 1030 nm. **b**, Left, fluorescence responses of GEVIs from a –70-mV baseline in voltage-clamped HEK293 cells. *n* = 5 (ASAP5) or 7 (ASAP7y) cells. Error bar, standard error of the mean (SEM). Gray area, physiological voltage range. Right, slopes of the above curves. **c**, Example fluorescence responses of GEVIs to current-evoked APs in cultured cortical neurons. To the right of each trace is an average of the measured electrical and optical AP waveform**s** induced by the injected current steps. Shaded area, SEM; *n* = 7 cells for each GEVI. **d,** Fluorescence changes from baseline during the plateau potential evoked by current injection; *n* = 7 cells for each GEVI. **e**, 1030-nm 500-fps scanless two-photon microscopy in organotypic hippocampal slice. Left, experimental setup and confocal image of ASAP7y-expressing neurons in slice. Right, example whole-cell current clamp and ASAP7y scanless two-photon voltage recordings. **f,** Fluorescence responses of ASAP7y to voltage changes imaged by 1000-nm two-photon resonant-scanning microscopy during whole-cell patch-clamping in mouse visual cortex *in vivo*; *n* = 3 cells. Error bar, SEM. **g**, Imaging of ASAP7y in mouse visual cortex by 1030-nm SLAP2 microscopy. Left, experimental schematic and image of an ASAP7y-expressing layer-2/3 neuron with scanned region indicated. Right, example trials of ASAP7y responses to gratings of different orientations presented in random order. Direction-specific responses comprise both APs and stimulus-locked subthreshold potentials (dashed box). **h,** Transcranial mesoscopy of ASAP7y with simultaneous LFP electrode recording. Left, raw image of ASAP7y in excitatory neurons after transduction by brain-penetrant AAV. Right, example LFP and ASAP7y traces. The electrode is placed near region 1. Regions 1 and 2 are located symmetrically across the midline. **i,** TEMPO fiber photometry of delta and gamma oscillations with ASAP7y. Left, example traces of PV interneurons in mouse cortex expressing ASAP7y or ASAP3 showing anesthesia-induced up and down states forming delta oscillations. Right, average up-state waveforms with timings determined by delta wave peaks. Shading, 95% confidence interval. **j,** Left, example ASAP7y and ASAP3 traces (colored) showing upward and downward state transitions in PV interneurons. Gray, traces filtered for the gamma (30-60 Hz) frequency band oscillations specifically during up-states. Right, plots of the mean fluorescence signal of ASAP7 or ASAP3 in the delta frequency band (1 ± 0.5 Hz, dashed traces; left axis), overlaid with plots of the delta phase-locked signal magnitude in the gamma band (30–60 Hz, solid traces; right axis). The convention used for the phase of delta oscillation is that 0-degree refers to the trough of the oscillation, i.e., the greatest hyperpolarization in the TEMPO signal. Shading, 95% confidence interval.

We compared ASAP7y to ASAP5 in cultured cortical neurons for brightness and responsiveness to natural voltage events. ASAP7y is brighter than ASAP5 with 490-nm one-photon excitation and as bright with 940-nm two-photon excitation, despite both conditions being optimized for the GFP-containing ASAP5. We then performed current injections while simultaneously recording evoked plateau potentials and action potentials (APs) electrically and optically. ASAP7y responded to APs with 61% larger changes than ASAP5, while responses to subthreshold activities were 140% larger (Fig. 1c-d). Thus, ASAP7y has higher SNR for both APs and subthreshold activity than ASAP5 when excited at typical wavelengths used for GFP-based indicators.

### ASAP7y improves subthreshold voltage imaging *in vivo* across imaging modalities

We examined if ASAP7y can be excited with Yb-doped fiber lasers of wavelengths of 1030–1040 nm. These high-power lasers have recently enabled two-photon imaging methods utilizing excitation beam parallelization for high sampling speeds, such as scanless two-photon microscopy and SLAP2 microscopy^47,48^. First, we characterized ASAP7y using scanless camera-based two-photon microscopy with a 1030-nm laser. Scanless two-photon microscopy of ASAP7y, combined with simultaneous whole cell patch-clamp recordings, detected spontaneous APs in neurons in organotypic hippocampal slices with high SNR (Fig. 1e).

We next tested ASAP7y’s responses to subthreshold depolarizations in mice. Simultaneous patch-clamp electrophysiology and 1000-nm raster-scanning two-photon imaging confirmed that ASAP7y has a steep response curve near resting membrane voltage *in vivo* (1.9% / mV; Fig. 1f). To verify the utility of ASAP7y for high-throughput *in vivo* microscopy using 1030-nm excitation, we tested ASAP7y using SLAP2 microscopy in visual cortex^47^ (Fig. 1g). In response to visual stimuli, ASAP7y detected both direction-selective subthreshold and spiking activity in single cells (Fig. 1g).

Given ASAP7y’s high responsivity to subthreshold voltage dynamics, we investigated whether ASAP7y population signals can mimic local field potential (LFP) measurements, which reflect predominantly subthreshold activity. We expressed ASAP7y broadly in excitatory neurons and imaged fluorescence noninvasively through the skull. ASAP7y signals near the site of an implanted electrode closely correlated with the LFP (r^2^ = 0.8; Fig. 1h; Extended Data Figure 2), exceeding previously obtained values for population voltage imaging^49^. A symmetrically located contralateral location displayed similar ASAP7y dynamics, as expected from the bilaterally symmetric nature of cortical circuit activation, while ASAP7y dynamics in a different cortical region exhibited lower coherence, supporting the specificity of the ASAP7y signal (Fig. 1h; Extended Data Figure 2).

A recent study found that ASAP3 population signals could track gamma oscillations using the TEMPO (transmembrane electrical measurements performed optically) approach, in which correction for motion and hemodynamic artifacts using a reference channel allows sensitive detection of low-amplitude oscillations^50^. We thus also tested ASAP7y in fiberoptic TEMPO experiments. ASAP7y reported anesthesia-induced up and down states in visual cortex that oscillated at delta frequencies, with 8-fold larger responses than ASAP3 (Fig. 1i), and detected gamma frequencies during up states with near 10-fold higher power at the same excitation irradiance (Fig. 1j). Compared to ASAP3, ASAP7y also detected gamma oscillations induced by visual stimuli with higher power and showed better photostability and less susceptibility to violet-light photoactivation (Extended Data Figure 3).

Taken together, these results show that ASAP7y allows reliable tracking of subthreshold voltage dynamics in the mammalian brain using a wide variety of imaging modalities, including those using high-power 1030-nm lasers.

### ASAP7y enables high-fidelity recording of voltage dynamics in *Drosophila*

Given our interest in examining how changes in voltage flow over the surface of individual neurons, we compared the properties of ASAP5 and ASAP7y in *Drosophila*, leveraging a well-characterized, non-spiking visual interneuron, Mi1 (Fig. 2a). Mi1 responds to increases in light, contrast increments, by depolarizing, and to decreases in light, contrast decrements, by hyperpolarizing^14,51^.We expressed ASAP7y and ASAP5 in Mi1, and presented animals with 8ms contrast increments and decrements. To achieve high sampling rates, we used acousto-optic deflector microscopy (AOD) to sample the Mi1 terminal in the M10 layer at 1041 Hz and recorded visually evoked responses at 940 nm or 1030 nm excitation. At 940nm, the peak excitation of ASAP5, we observed detectable single trial responses from ASAP5-expressing Mi1 terminals (9.7 ± 1.6% peak ΔF/F), and substantially larger responses from ASAP7y-expressing Mi1 terminals (20.3 ± 3.9% peak ΔF/F) (Fig. 2b-d). At 1030nm, a wavelength that is not compatible with ASAP5 excitation, ASAP7y performance was comparable to that seen at 940nm (22.3 ± 2.9% peak ΔF/F, Fig. 2b-d). ASAP7y also displayed improved brightness over ASAP5 at 940nm and remained bright at 1030nm (Fig. 2e). As a result, the peak SNR of ASAP7y was ∼2.3-fold better than ASAP5 (Fig 2f). Finally, we also observed that ASAP7y displayed significantly better photo-stability than ASAP5 (Fig. 2g).

**Figure 2.**
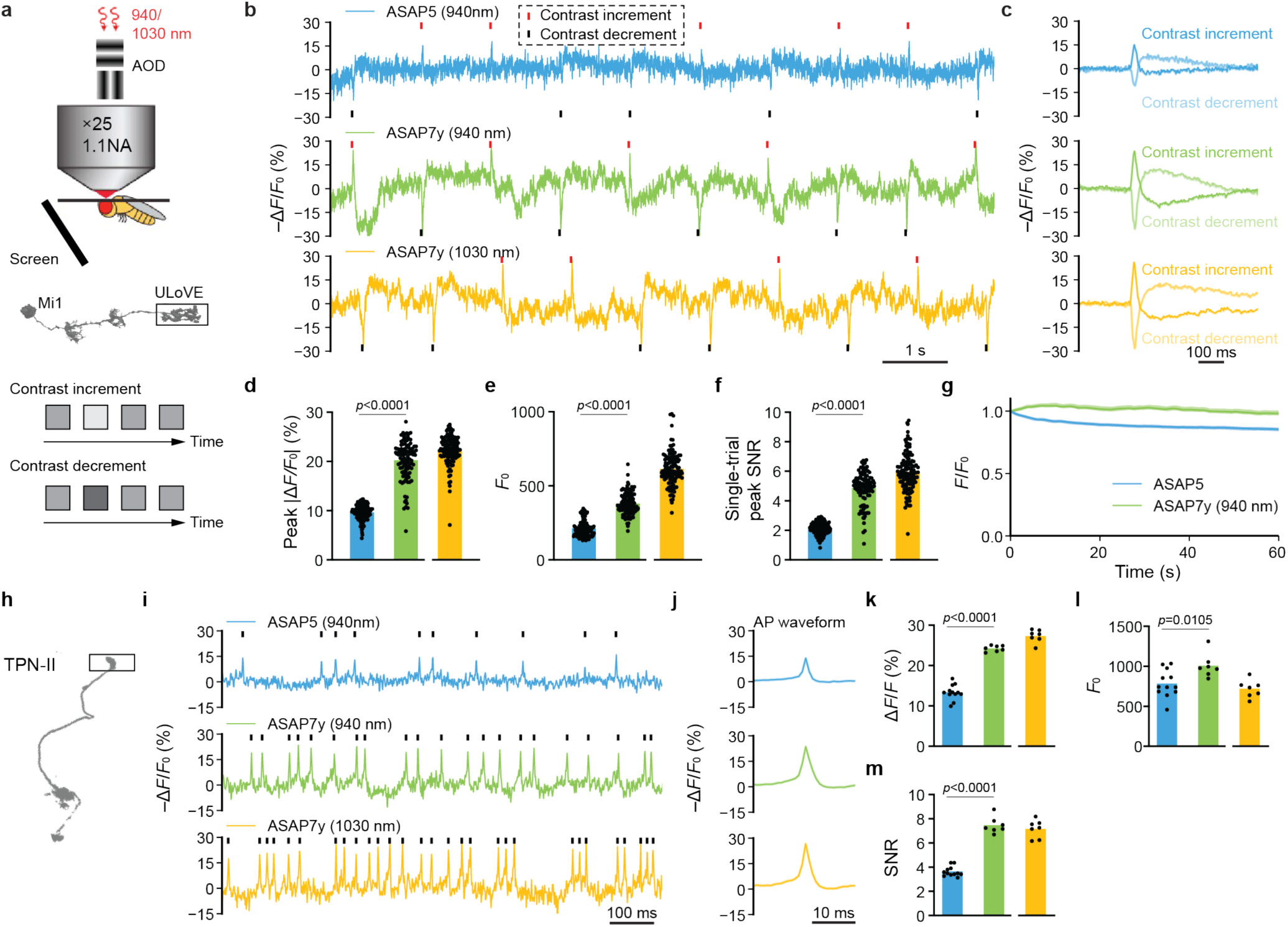
ASAP7y exhibits improved SNR and photo-stability for two-photon imaging *in vivo*. **a**, Imaging visually evoked responses in Mi1neurons *in vivo* using ASAP5 and ASAP7y. Imaging was performed using an AOD random access microscope combined with 940 nm or 1030 nm excitation. Stimuli were contrast increments or decrements of 8ms duration. **b**, Example traces from individual Mi1 neurons expressing ASAP5 or ASAP7y evoked by increments (red ticks) and decrements (purple ticks). **c**, Stimulus triggered averages (STA) from single cells corresponding to traces in (**b**) (*n* = 30 trials for each stimulus). Shaded area, standard error of the mean (SEM). **d-f**, Quantification of peak responses, baseline fluorescence and peak SNR in Mi1 neurons. Each dot is a neuron. Peak SNR was calculated based on single trials; *n=*108 cells, 6 flies for ASAP5 (940 nm), *n =* 108 cells, 6 flies for ASAP7y (940 nm), *n =* 124 cells, 6 flies for ASAP7y (1030 nm). Differences between ASAP5 and ASAP7y at 940 nm for peak ΔF/F (\*\*\**P* <0.0001), brightness (\*\*\**P* <0.0001), and peak SNR (\*\*\**P* <0.0001) were all significant as quantified by two-tailed *t*-test. We did not compare measurements of ASAP7y across wavelengths. **g**, Normalized fluorescence of ASAP5 and ASAP7y over 1 min of continuous imaging in Mi1. Shaded area, SEM. **h**, Imaging spontaneous spiking activity in TPN-II neurons expressing ASAP5 or ASAP7y under 940 nm or 1030 nm excitation. **i**, Example recordings of individual TPN-II neurons. **j**, Average waveform of APs corresponding to traces in (**i**). Shaded area, SEM; *n* = 1084 APs for ASAP5 (940nm), 949 APs for ASAP7y (940nm) and 738 APs for ASAP7y (1030nm). **k-m**: Quantification of peak responses, baseline fluorescence and peak SNR of TPN-II neurons imaged. *n* = 12 cells, 6 flies for ASAP5 (940 nm), *n* = 7 cells from 4 flies for ASAP7y (940 nm), *n =* 7 cells from 4 flies for ASAP7y (1030 nm). Differences between ASAP5 and ASAP7y at 940nm for peak response (\*\*\**P* < 0.0001), brightness (\**P* = 0.0105), and SNR (\*\*\**P* < 0.0001) were all significant as quantified by two-tailed *t*-test.

Next, we compared the performance of ASAP7y and ASAP5 in reporting APs *in vivo*, leveraging TPN-II, a thermosensory neuron that fires APs spontaneously at room temperature (Fig. 2h)^52,53^. We sampled TPN-II neurons expressing either ASAP7y or ASAP5 at 1041Hz using both 940 nm and 1030 nm excitation and recorded spontaneous APs. ASAP7y exhibited a peak response to APs that was 81% higher than ASAP5, displayed increased brightness, and had a two-fold better SNR at 940nm (Fig. 2i-m). Moreover, ASAP7y exhibited comparable peak responses at 1030 nm excitation (Fig. 2i, k). Strikingly, due to the increased photostability of ASAP7y, we could routinely record APs using 1030nm excitation of TPN-II neurons for 20 minutes (Extended Data Figure 4).

### *In vivo* measurements of electrical conduction in single neurons reveal heterogeneous subcellular voltage distributions

To gain insight into how changes in membrane potential propagate along individual neurons *in vivo*, we sought to capture single-trial, multi-site voltage responses with millivolt precision and kilohertz resolution across a diversity of cell types in the fly visual system (Fig 3a). We used two-photon acousto-optic deflector-based random-access microscopy, in combination with expression of ASAP7y in single cells of genetically defined types ^4,40,54^. To capture a broad spectrum of neuronal diversity, we selected cell types that varied in function, molecular identity, and morphology, projected across multiple visual ganglia, and ranged in size from tens to hundreds of microns. Following receptive field mapping (Fig. 3b, Extended Data Figure 5), we presented a minimal stimulus - a single dark spot (r = 4°) - for 64 ms at the receptive field center and recorded membrane potential at multiple regions of interest (ROIs) along the neuron (Fig. 3c-t, Extended Data Figure 6).

**Figure 3.**
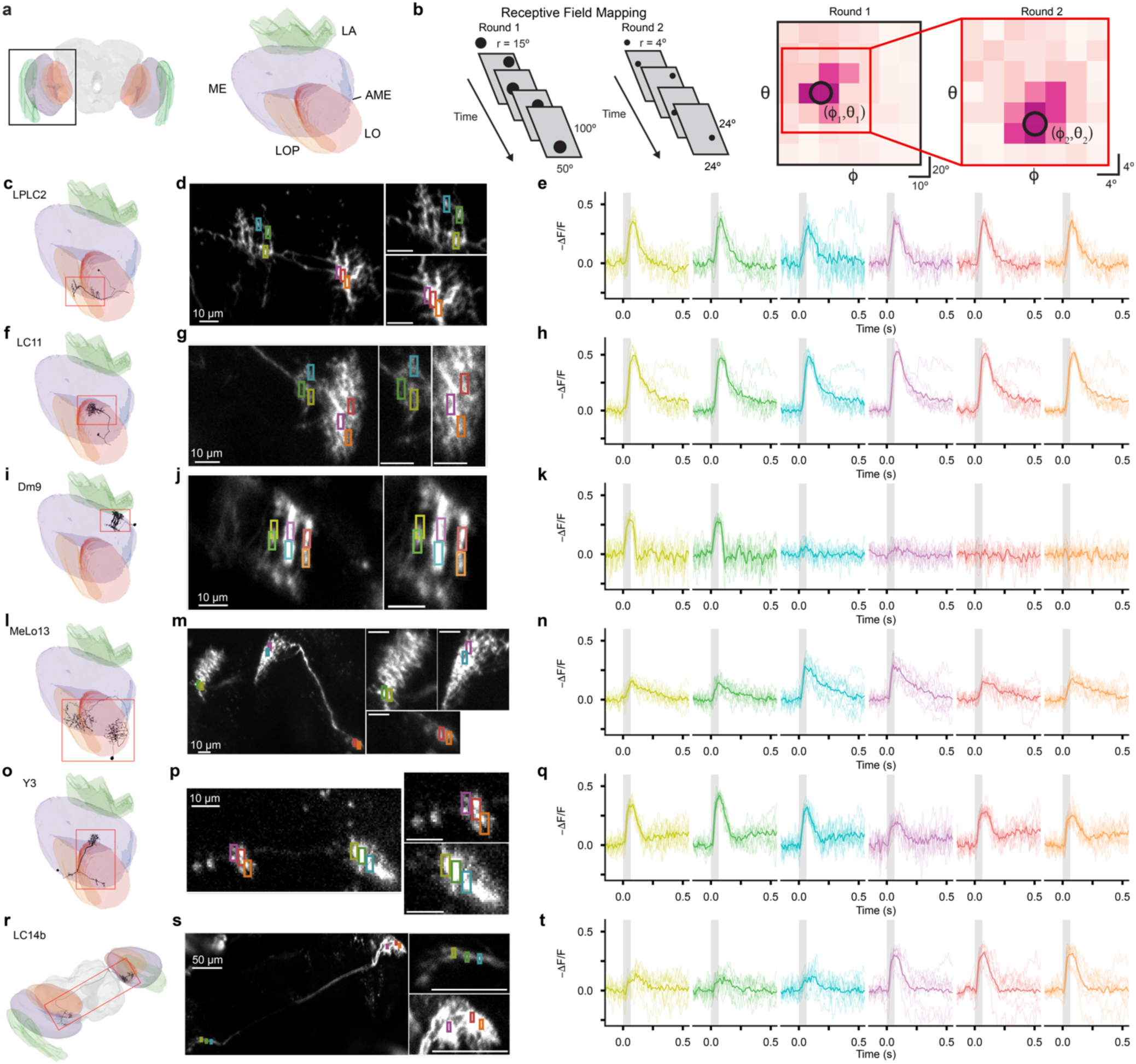
ASAP7y enables single trial measurements of visually evoked responses in subcellular domains. **a**, Frontal view of the *Drosophila* brain. Inset, right optic lobe with each ganglion labelled: Lamina (LA, green), Medulla (ME, purple), Accessory Medulla (AME, blue), Lobula (LO, orange), and Lobula Plate (LOP, yellow). **b,** Schematic of iterative receptive field mapping. In the first round, a dark spot (r = 15°) was presented on a uniform grey background across a coarse spatial grid (φ = –30° to 20°; θ = –50° to 50°, 10° steps). In the second round, a small, dark spot (r = 4°) was presented across a 24°-by-24° grid centered on the peak response location from the first round (φ = –12° to 12°; θ = –12° to 12°, 4° steps). Responses were quantified as mean −Δ*F*/*F* within 150 ms following stimulus onset, and a 2D Gaussian was fit to these responses to estimate the receptive field center. **c,** Reconstruction of a portion of a single LPLC2 neuron in the optic lobe. The red box denotes the imaging window. **d,** A single LPLC2 neuron, with 2 × 6 *um* regions of interest (ROIs) displayed **e,** −Δ*F*/*F* from the ROIs shown in (**d**) over 10 stimulus trials, in response to a small, dark spot stimulus (r = 4°, duration = 64 ms) presented at the receptive field center on an otherwise uniform grey background. Individual trials are shown as thin lines; the mean response over 10 trials is shown in bold; grey shading denotes the stimulus presentation window. **f-h**, same as (**c-e**) for LC11. **i-k,** same as (**c-e**) for Dm9. **l-n**, same as (**c-e**) for MeLo13. **o-q**, same as (**c-e**) for Y3. **r-t**, same as (**c-e**) for LC14b.

We first examined LPLC2, a lobula plate/lobula columnar (LPLC) visual projection neuron previously characterized as selective for looming stimuli (Fig. 3c-e) ^27,55,56^. LPLC2 displayed a monophasic depolarization that extended for ∼200 ms in response to the stimulus, with near-uniform depolarizations across ROIs placed on neurites in the lobula and the lobula plate (Fig. 3d, e). We next examined LC11, a lobular columnar (LC) neuron that responds preferentially to small moving objects^55^. Similarly to LPLC2, LC11 exhibited a stimulus-evoked monophasic depolarization, with near-uniform voltage responses from ROIs placed in different layers of the lobula (Fig. 3f-h). Comparable results were obtained from LC4, a loom-sensitive LC neuron (Extended Data Figure 6)^55,56^.

We next imaged Dm9, a small, multi-columnar interneuron in the medulla implicated in color opponency^28^ (Fig. 3i-k). Dm9 responded robustly to stimuli with a monophasic depolarization lasting ∼100ms (Fig. 3k). However, unlike LPLC2, LC11, and LC4, Dm9 exhibited localized voltage responses: ROIs placed along the same columnar branch displayed strong responses, while ROIs placed along neighboring branches exhibited minimal responses (Fig. 3j, k), indicating that changes in voltage do not propagate uniformly across retinotopically organized columns. Comparable results were obtained from another highly branched neuron, CT1 (Extended Data Figure 6).

We observed non-uniform voltage distributions across the arbors of three more cell types. MeLo13, a previously uncharacterized neuron with arbors in both the lobula and the medulla (Fig. 3l-n), responded to the stimulus with monophasic depolarizations that had different amplitudes across distant projections in the medulla and lobula, and at the soma (Fig. 3m, n). Similar differences in voltage amplitudes were seen across the medulla and lobula projections of Y3, a neuron implicated in motion processing^57^ (Fig. 3o-q). LC14b, a bilaterally projecting LC neuron previously implicated in binocular comparisons, also exhibited non-uniform voltage responses between ipsi- and contralateral arbors (Fig. 3r-t)^58^.

Taken together, these results demonstrate that ASAP7y, combined with a cell-type selective genetic driver, enables routine kilohertz sampling of subcellular voltage signals across individual neurons *in vivo*. This approach revealed strikingly diverse subcellular voltage distributions across cell types. Some neurons, namely LPLC2, LC11 and LC4, exhibited near-uniform voltage responses across all of their measured projections. By contrast, other cell types, including Dm9, MeLo13, Y3 and LC14b, displayed marked differences in their voltage distributions across spatial scales ranging from tens to hundreds of microns. These results raise the possibility that there is a diversity of electrical properties across cell types, ranging from electrically compact neurons with uniform voltage distributions to compartmentalized neurons with localized voltage distributions.

### Diverse morphological features drive heterogeneity of voltage distribution across cell types

How might this heterogeneity in subcellular voltage distributions emerge? One potential mechanism is rooted in cell-type specific differences in neuronal morphology. Leveraging the nanometer-resolution reconstruction of the visual system^4^, we simulated voltage propagation in individual neurons to test this possibility. We built morphologically constrained models with passive membrane properties for each neuron in the visual system, injected current at specific sites, and recorded voltage dynamics across each *in silico* cell (Fig. 4a, b; Extended Data Figure 7). To capture the maximal physiologically relevant voltage decay, we chose the injection site as the location with the highest density of synaptic inputs from the dominant pre-synaptic partner, and we analyzed the output synapse with the lowest voltage signal as the output site.

**Figure 4.**
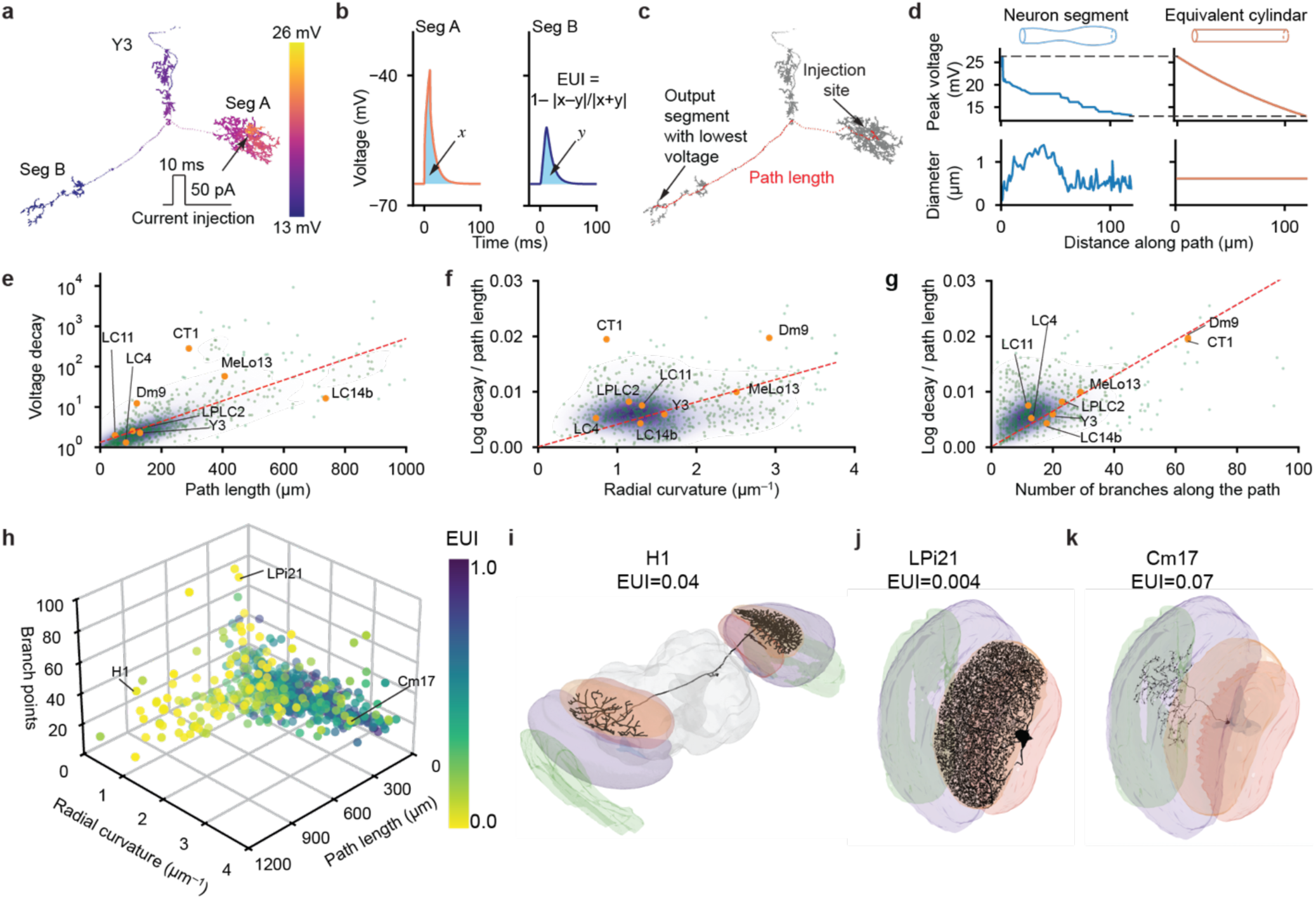
Morphological features tune cell-type specific electrotonic properties. **a**, Example voltage distribution across a simulated Y3 neuron. Schematic of current injection site (arrow) and waveform (trace). **b**, Averaged voltage dynamics of segment A and segment B from the simulation in (**a**). EUI was computed on the area under the voltage dynamics curves. **c**, Identifying the path from injection site to the output segment with the lowest voltage. **d**, Computing the radial curvature of the equivalent cylinder of a neurite with varying radius. **e**, Distribution of median path lengths and median voltage decay across individual simulated neurons of each type. Orange dots highlight the neuron types we measured experimentally. Color shade (purple) denotes density of dots. **f**, Distribution of the median logarithm of voltage decay and the median radial curvature across individual simulated neurons of each type. Color shade (purple) denotes density of dots **g**, Distribution of the median logarithm of voltage decay and median number of branches across individual simulated neurons of each type. Color shade (purple) denotes density of dots **h**, Distribution of neuron types in geometric feature space: path length (in μm), equivalent radial curvature (in μm^-1^) and number of branch points along the path. Each dot is colored by the median EUI of the type. Three neuron types with extreme geometric features are highlighted in **i-k**.

Next, we quantified the morphological parameters that shape voltage decay (Fig. 4a-d). Cable theory predicts that voltage decays exponentially along an infinitely long cylinder at steady state^10^. Based on this intuition, we first identified the path length from the injection site to the output site (Fig. 4c), computed the voltage decay for each individual neuron, and plotted the median decay and path length for each neuron type (Fig. 4e). We observed that the simulated voltage decay and the path length were highly correlated across neuron types over more than two orders of magnitude, indicating that path length contributes significantly to voltage decay (Fig. 4e; Pearson correlation coefficient (PCC) = 0.82). Intriguingly, we observed that neurons with short path lengths could still achieve large voltage decays comparable to those seen in neurons with much longer path lengths, suggesting that additional morphological parameters might be leveraged to shape voltage decay. To explore these additional factors, we first computed the logarithm of the voltage change to account for its exponential decay and normalized this quantity by the path length. As the space constant of the decay function in an idealized cable is influenced by its radial curvature, we next computed the equivalent cylinder, the cylinder with uniform radius that has the same path length and voltage decay as the neuron, for each individual neuron (Fig. 4d). We then plotted the median normalized voltage decay against the median radial curvature of each neuron type (Fig. 4f). Radial curvature exhibited a significant correlation with the normalized voltage decay (PCC = 0.27), indicating that across these neuron types, radial curvature is used to tune voltage decay. In a similar vein, we next examined the relationship between branch number and normalized voltage decay. We counted the number of branches along the path from the input site to the output site of each neuron and plotted the median of the branch number and the median of the normalized voltage decay for each neuron type (Fig. 4g). Branch number was highly correlated with the normalized voltage decay (PCC = 0.57), revealing that branching is also used to shape the voltage transform across these neurons.

Taken together, across these cell types, there are at least three geometric features that tune the physiologically relevant voltage transformation: the path length, the radial curvature of the neurite, and the number of branches along the path. To quantify the degree of voltage decay across each neuron, we developed an Electrical Uniformity Index (EUI) (Fig. 4b). This metric has a range from 0 to 1, with 1 indicating an iso-potential voltage distribution with no decay, and 0 indicating a fully compartmentalized, localized voltage distribution, with complete decay. To examine how different cell types might combine morphological features to shape voltage transformations, we plotted the joint distribution of path length, radial curvature and branch number, and colored each cell type by its median EUI (Fig 4h). This distribution was highly skewed, with prominent tails along each of the three axes, with significant compartmentalization at the extrema of all three factors. That is, comparable compartmentalization can be achieved by a long path length, as exhibited by the bilaterally-projecting H1 neuron (Fig. 4i), by a highly branched structure, as exhibited by the interneuron LPi21 (Fig. 4j), or by very thin neurites, as exhibited by the interneuron Cm17 (Fig. 4k). This skewed distribution suggests that individual cell-types preferentially leverage a specific subset of morphological parameters to tune their voltage transform, perhaps reflecting physiological or connectomic constraints.

### Quantification of *in vivo* voltage dynamics reveals cell-type specific organization of electrical signals

To quantify cell-type specific voltage dynamics, we measured voltage dynamics within and between recognizable morphological features in individual cells spanning eight cell types (Fig. 5a, b). Regions were chosen with the goal of interrogating the relationship between distinct subcellular compartments. Depending on neuronal morphology this meant sampling projections across different ganglia, isolating individual columnar branches, or contrasting dendritic and axonal domains in polarized neurons (Fig. 5b, Extended Data Figure 8). Importantly, voltage responses within a region were homogenous (Extended Data Figure 9), enabling robust regional comparisons across individual cells of a given cell type. To characterize voltage dynamics across a range of stimulus intensities, we presented a single dark spot (r = 4°) at the receptive field center across four durations (8, 16, 32, and 64 ms) (Fig. 5c). Each stimulus was presented ten times, and for each trial, we integrated the voltage response across regions (Fig. 5c). To quantify the relationship between the voltage signals in each region of each neuron, we computed the EUI from experimental measurements of each trial and fit a linear regression for each individual neuron across all trials (Fig. 5d).

**Figure 5.**
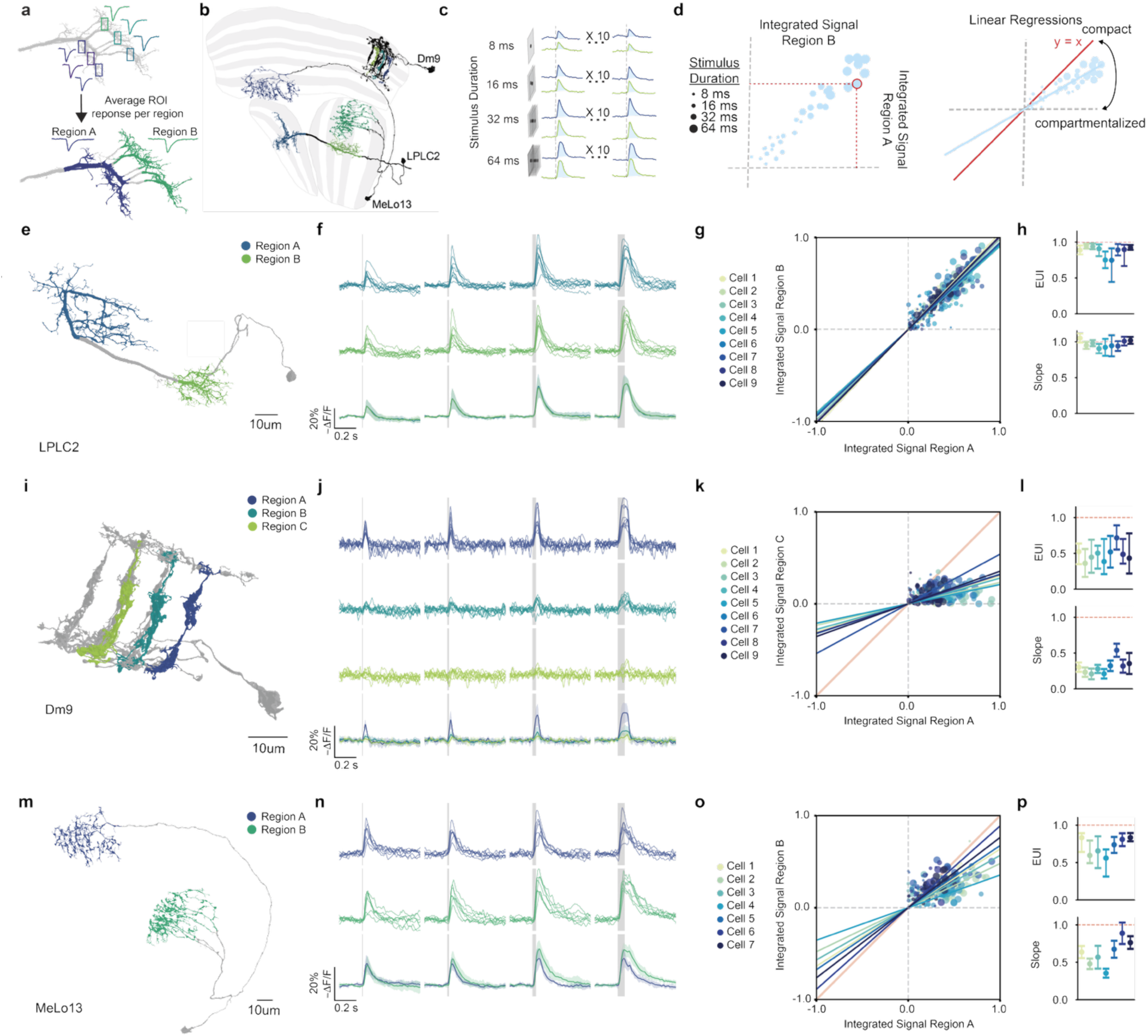
*In vivo* measurements of electrical conduction in single neurons reveal heterogeneous subcellular voltage distributions. **a,** Schematic illustration of ROI placement and region definition. **b**, Schematic of three imaged neurons (LPLC2, MeLo13, and Dm9) with measured regions highlighted, overlaid on the optic lobe. **c**, Flies were presented with a small, dark spot of radius = 4° for varying durations (8, 16, 32, and 64 ms), randomly interleaved, with each stimulus repeated across 10 trials. For each region, trial-wise responses were quantified by integrating the –Δ*F*/*F* trace over a defined time window to obtain an area-under-the-curve (AUC) metric. **d**, Left: Example plot of the integrated signal in region A vs. Region B for a single neuron across 40 trials. Each point represents a single trial (4 stimulus conditions × 10 presentations each = 40 trials). Calculation of the EUI for a single trial is indicated by the red dashed lines, and is calculated as: 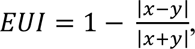 integrated signals in Regions A and B. Right: Schematic of the zero-intercept linear regression performed per neuron across all stimulus trials. A slope approaching 1 (*y = x*, denoted in red) indicates an electrically compact neuron, with a uniform voltage distribution across regions, while a slope approaching 0 indicates an electrically compartmentalized neuron, with non-uniform, localized voltage distributions across regions. **e**, Morphological reconstruction of an LPLC2 neuron with regions A and B indicated. **f**, Stimulus-evoked responses of measured LPLC2 neurons (*n* = 9 cells, 8 flies) across stimulus durations. Stimuli are shown from left to right in order of increasing duration (8, 16, 32, and 64 ms), as indicated by grey shading. Top: Each line represents the mean –Δ*F*/*F* in region A for a recorded LPLC2 neuron over 10 stimulus presentations. Middle: Corresponding mean –Δ*F*/*F* in Region B. Bottom: Overlay of mean responses in region A and region B across all recorded LPLC2 neurons. Shading, standard deviation in region A and region B. **g**, Integrated signal in Region B plotted against Region A over all measured LPLC2 neurons and all trials. Each color represents trials from an individual neuron. Data are plotted as described in (**d**), with zero-intercept linear regression fits overlaid for each cell. **h**, Quantification of EUI and slope of zero-intercept linear regressions across measured LPLC2 neurons. Top: quantification of EUI for each LPLC2 neuron over all trials. Data shown as median with 25^th^–75^th^ percentile error bars. Population median = 0.90. Bottom: quantification of constrained linear regressions for each LPLC2 neuron over all trials. Data shown as slope with the 95% confidence interval in error bars. Population median = 0.95. **i**–**l**, same as **e**–**h** for Dm9 regions A, B and C (*n* = 9 cells, 9 flies). **l**, Quantification of EUI and slope of zero-intercept linear regressions across measured Dm9 neurons (region A to region C). Top: population median = 0.49. Bottom: population median = 0.30. **m**–**p**, same as **e**–**h** for MeLo13 (*n* = 7 cells, 7 flies). **p**, Quantification of EUI and slope of zero intercept linear regressions across measured MeLo13 neurons. Top: population median = 0.74. Bottom: population median = 0.64. For quantification of all individual cells across all cell types see **Extended Data Table 1**.

We first examined LPLC2, focusing on neurites in the lobula and the lobula plate to measure voltage responses across visual ganglia (Fig. 5e-h). LPLC2 displayed near uniform integrated voltage signals across regions, with a median EUI of 0.90, and a median slope of 0.95 across cells (Fig. 5f-h). LC11 and LC4 also displayed near-uniform voltage responses across lobula layers (Extended Data Figure 10). Next, we imaged Dm9 (Fig. 5i-l). We measured voltage dynamics in individual branches, each corresponding to a single retinotopic column, and observed integrated voltage signals localized to each branch. Branches in adjacent columns displayed a median EUI of 0.55 and slope of 0.40 (Extended Data Figure 10), while branches in non-adjacent columns displayed a median EUI of 0.49 and a slope of 0.30 (Fig. 5j-l), suggesting that individual retinotopic columns in Dm9 are electrically compartmentalized. We next examined voltage dynamics in the medulla and lobula arbors of MeLo13 (Fig. 5m-p). Voltage responses differed between lobula and medulla arbors, with a median EUI of 0.74, and a slope of 0.64, indicating that these projections are also incompletely coupled (Fig. 5n-p). Voltage compartmentalization was also seen between projections in the medulla and lobula plate in Y3 and between contralateral projections in LC14b (Extended Data Figure 10).

Our results reveal cell-type specific differences in voltage dynamics. In some neurons, voltage distributions between measured neurites were uniform (LC4, LC11, LPLC2), a result that could emerge from either identical patterns of synaptic input to each neurite, or from unimpeded electrical conduction across the cell. To distinguish between these possibilities, we modeled voltage propagation between the measured regions of each of these cell types across physiologically plausible biophysical parameters (Extended Data Figure 11). These results support the notion that these cells are electrotonically compact. In other neurons, measured neurites displayed non-uniform voltage distributions (Dm9, MeLo13, LC14b, Y3), a result that can only emerge from some degree of electrotonic compartmentalization. In line with this conclusion, our modeling demonstrates that these neurons are compartmentalized across physiologically plausible biophysical parameters (Extended Data Figure 11). Taken together, these findings demonstrate that uniformity and compartmentalization represent the ends of a continuum of electrical organization that defines a fundamental axis of cell-type diversity.

### Electrotonic compartmentalization creates neural substrates for parallel processing in single neurons

We next investigated whether subcellular electrotonic domains within a neuron are functionally specialized. Given that Dm9 is a multi-columnar neuron that receives retinotopically organized input, we hypothesized that each branch might correspond to a different point in visual space. We therefore mapped the spatial receptive field of each branch and discovered that each had a unique receptive field center (Fig. 6a, b). In contrast, in electrotonically compact neurons (LC4, LC11, LPLC2), all ROIs on the same cell had the same spatial receptive field regardless of placement (Fig. 6c, d; Extended Data Figure 12). Cross correlation analysis revealed that Dm9 branches were tuned to distinct regions of visual space, while ROIs placed within the same branch were tuned to the same region of visual space (Fig. 6e). Conversely, electrotonically compact neurons exhibited no spatial displacement between ROIs (Fig. 6e). To examine how inputs to one branch are represented across its neighbors, we presented a single dark spot at the receptive field center of each branch and recorded the responses across all branches (Fig. 6f). Each branch exhibited the strongest voltage response when stimulated at its own receptive field center, while neighboring branches displayed much weaker responses (Fig. 6f). We next constructed a matrix that described the relative response amplitudes for each branch and generated a modeled matrix in which we simulated the voltage responses evoked by current injection into each branch (Fig. 6g; Extended Data Figure 13). Both of these matrices displayed crosstalk, suggesting that electrotonic propagation could account for the observed *in vivo* interactions between branches. We also imaged CT1, a large amacrine cell, and found that it too displays electrical compartmentalization between retinotopically organized branches (Fig. 6e, Extended Data Figure 14). This observation is consistent with previous work showing that visually evoked Ca^2+^ responses in CT1 are compartmentalized^24^. Taken together, Dm9 and CT1 exemplify how voltage compartmentalization enables a single neuron to represent multiple, spatially distinct input channels in parallel while also facilitating integration of information spanning multiple branches. In contrast, LC4, LC11, and LPLC2, despite also receiving retinotopically organized inputs across multiple columns, exemplify how electrotonic compactness enables neurons to uniformly pool information across visual space.

**Figure 6.**
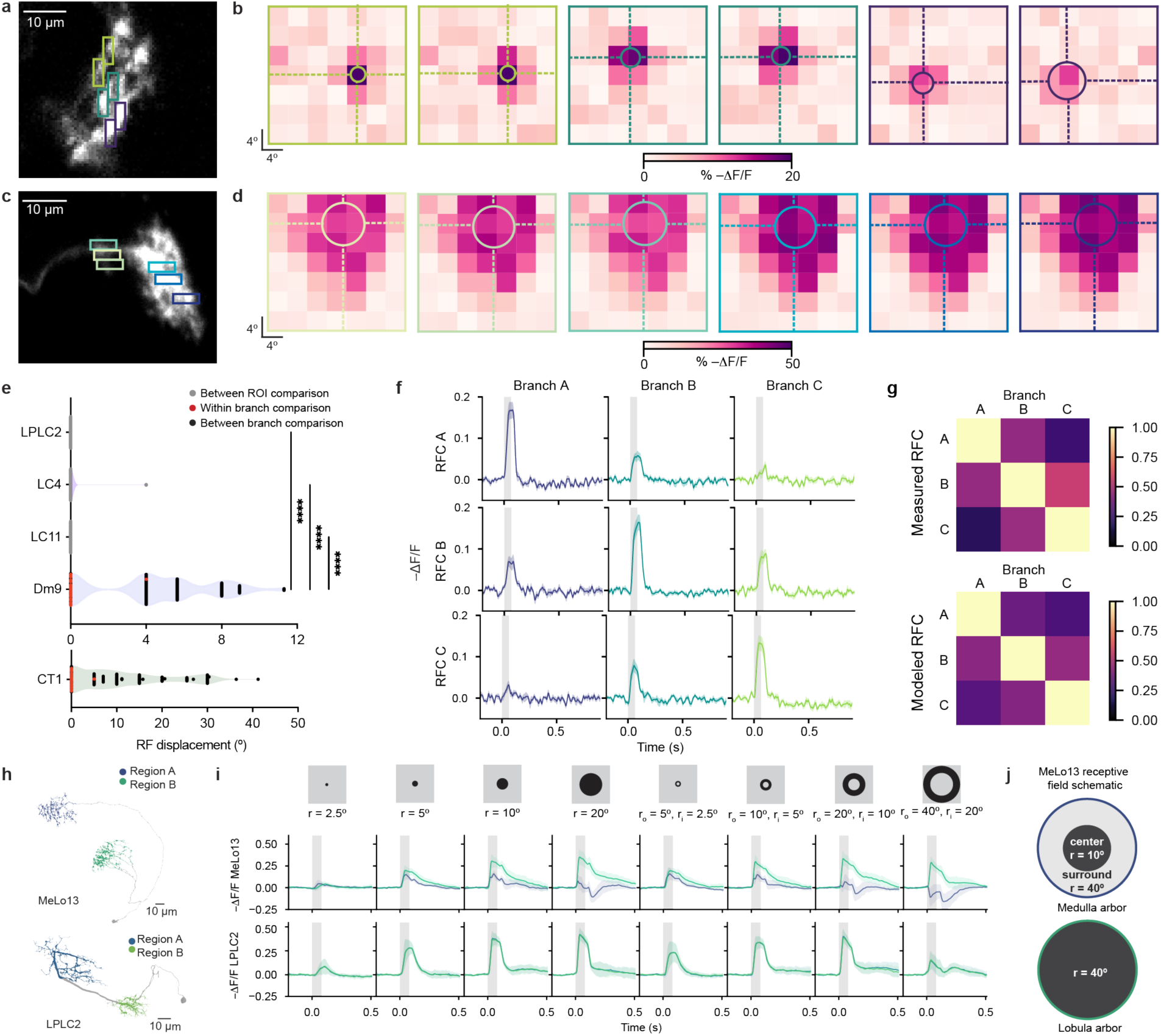
Voltage compartmentalization produces functionally specialized subdomains. **a**, Imaged Dm9 neuron with overlaid ROIs. **b,** Spatial tuning maps for each ROI. A 2D Gaussian was fit to –Δ*F*/*F* to extract the center and extent of the peak response, visualized as overlaid ellipses. **c, d,** Same as **a, b** for an imaged LC11 neuron. **e,** Receptive field shifts between ROIs within single neurons (6 ROIs per neuron, 15 pairwise comparisons) aggregated across all neurons of each cell type (LPLC2, *n* = 9; LC4, *n* = 8; LC11, *n* = 9; Dm9, *n* = 9; CT1, *n* = 7). Significant differences were observed between cell types (ordinary one-way ANOVA: *F* (4, 598) = 243.6, *\*\*\*\*P* < 0.0001). Post-hoc Tukey’s multiple comparisons test revealed significant differences between Dm9 and LPLC2, LC4, and LC11 (Dm9 vs. LPLC2, **** *P* < 0.0001; Dm9 vs. LC4, **** *P* < 0.0001, Dm9 vs. LC11, **** *P* < 0.0001) with no significant differences between LC11, LC4, and LPLC2 (LC11 vs. LC4, NS, *P* = 0.9710; LC11 vs. LPLC2, NS, *P* > 0.9999; LC4 vs. LPLC2, NS, *P* = 0.9660). CT1 receptive field mapping was performed over a broader 100°-by-50° stimulus grid with 10° resolution and was therefore excluded from statistical comparisons. **f**, Branch-specific Dm9 responses across all measured neurons (*n* = 9 cells, 9 flies). The receptive field center of each Dm9 branch was mapped, and a small dark-spot stimulus (r = 4°) was presented at the receptive field center of each branch. Responses were recorded across all branches, and the average response per branch was calculated for each neuron. Data are shown as the mean –Δ*F*/*F* per branch across Dm9 neurons. Colored shading, SEM. **g**, Measured (top) and modeled (bottom) confusion matrices. Pixel (*i*, *j*) is the integrated signal of branch *i* to the stimulation of branch *j*. **h**, Morphological reconstructions of MeLo13 and LPLC2 neurons with regions highlighted (as in Fig. 5e, m). **i**, Schematic of stimulus set comprising dark spots and annuli of increasing radii. Data shown as mean –Δ*F*/*F* across MeLo13 neurons (*n* = 7 cells, 7 flies) or across LPLC2 neurons (*n* = 9 cells, 8 flies) for regions A and B. Regions are color-coded as in (**h**). Colored shading, SEM; grey shading, stimulus duration. **j**, Schematic illustration of the receptive field structure in the medulla arbor (Region A) and the lobula arbor (Region B) of MeLo13 neurons.

We next wondered whether expanding our stimulus regime would reveal additional diversity in receptive field organization across subcellular compartments. Given that all cell types in our dataset had OFF responses, we presented a series of dark dots and annuli of varying diameters to explore spatial receptive field structure across subcellular regions of individual neurons (Fig. 6h-i; Extended Data Figure 15). Strikingly, we found that MeLo13 displayed unique spatial receptive fields in different subcellular regions, even displaying simultaneous hyperpolarizing and depolarizing responses in different regions (Fig. 6i). Moreover, the medulla arbor of Melo13 exhibited a clear center-surround organization, while the lobula arbor of MeLo13 lacked opponent structure, and instead had a large OFF receptive field (Fig. 6j). In contrast, electrotonically compact neurons shared common spatial receptive field organizations across subcellular regions (Fig. 6h, i; Extended Data Figure 15). Thus, the divergence of spatial receptive field structures across subcellular compartments of MeLo13 provides another example in which compartmentalization supports parallel computations within a single cell.

Taken together, our results reveal how electrotonic properties shape neuronal feature selectivity. Electrotonically compact neurons have uniform voltage distributions and therefore pool information from inputs anywhere on their arbor, integrating over time. Electrotonically compartmentalized neurons have non-uniform voltage distributions and can therefore integrate over both their complex morphology and over time. As a result, compartmentalized neurons can simultaneously process information within and across electrical domains in parallel.

## Discussion

### Summary

Understanding how changes in membrane potential are re-shaped as they propagate from inputs to outputs is fundamental to a mechanistic understanding of single neuron computation. However, measuring such changes in membrane potential with subcellular resolution and sufficient temporal resolution has remained challenging *in vivo*. As a result, the fundamental question of whether and how different cell types might organize these signals across subcellular domains to implement functionally important computations has remained incompletely understood. Here we develop ASAP7y, a novel GEVI with unprecedented subthreshold responsivity that facilitates sensitive and extended sampling of *in vivo* voltage dynamics. We combine ASAP7y with random-access two-photon microscopy and sparse labeling in genetically defined cell types to simultaneously record sensory stimulus-evoked voltage dynamics with millivolt and millisecond precision at multiple sites along arbors of individual neurons.

These recordings reveal cell type specific stimulus-evoked voltage distributions, with some cell types displaying nearly uniform changes in membrane potential across all neurites, while other cell types display non-uniform changes in membrane potential that define sub-cellular electrical domains. Leveraging a biophysical simulation of more than seven hundred cell types, we reveal that specific geometrical parameters of neurites, namely path length, branch number and radial curvature, predict a vast continuum of cell-type specific electrotonic properties, illustrating how diverse voltage distributions can arise from underlying morphology. Finally, we show that different subcellular domains within a single neuron can have distinct feature selectivity, demonstrating that electrotonic compartmentalization creates substrates for functionally important local computations. As a result, cell-type specific tuning of voltage propagation greatly expands the computational capacity of single neurons.

### ASAP7y enables *in vivo* single trial measurements of subthreshold changes in membrane potential

ASAP7y displays unprecedented responsivity to small voltage fluctuations. Compared to ASAP5, one of the class-leading sensors of sub-threshold voltage potentials, ASAP7y is more than twice as responsive at resting membrane potentials (2.7% vs. 1.3% per mV), is brighter, and exhibits wider two-photon excitation compatibility. As we demonstrate here, this combination of properties makes it possible to record subthreshold changes in membrane potential, in multiple subcellular domains of single neurons, with high temporal resolution *in vivo*. Furthermore, as a result of its red-shifted excitation spectrum, ASAP7y can be combined with RFP-based sensors or blue-shifted opsins, opening new opportunities for multiplexed recordings and perturbations. The red-shifted excitation of ASAP7y also enables the use of high-power Ytterbium-doped fiber lasers as excitation sources, facilitating many high-throughput two-photon microscopy methods^16,47,48^. Finally, this sensor works well when expressed in rodent cortex, setting the stage for parallel studies of subcellular voltage compartmentalization across species.

### Electrotonic properties are regulated cell-type specifically

Prior work has left unclear the extent to which neurons display voltage compartmentalization *in vivo^2,3,59^*. Our measurements of voltage dynamics in many cell types revealed that different cell types employ distinct modes of electrical signal distribution, ranging from uniform to highly localized distributions. These observations expand on a fundamental axis of cell type diversity that shapes neuronal function^60^. *In vivo* measurements of voltage dynamics across subcellular regions inherently contain information about both the spatiotemporal pattern of synaptic input and voltage propagation. For some compartmentalized neurons, we observed stimulus-dependent changes in voltage distributions, highlighting the richness of the interactions between inputs and voltage propagation and emphasizing the importance of studying single neuron computation *in vivo*. Finally, our approach of measuring subcellular changes in membrane potential at high temporal resolution in individual cells is applicable across cell types and model organisms, enabling voltage dynamics to be explored in many contexts.

This novel axis of cell type diversity emerges in part from neuronal morphology, which has recently been characterized at nanoscale resolution through connectomics^4,41^. We used nanoscale EM reconstructions of each cell in combination with biophysical modeling to explore the electrotonic properties of 717 cell types in the visual system. As expected from prior theoretical work, these simulations revealed that path length, the radial curvature of the neurite, and the number of branches along the input-output path are key geometrical parameters that shape voltage propagation. Intriguingly, however, our analysis revealed that individual cell types predominantly weight particular combinations of these features to tune their voltage propagation, yielding a highly skewed distribution. For example, compartmentalization can emerge through long neurites, thin neurites or highly branched neurites, morphological features that are rarely combined at their extrema in this dataset. We infer that additional developmental, biophysical and connectomic constraints play central roles in dictating the cellular mechanisms that underpin voltage compartmentalization in specific cell types.

While our simulations correctly classified neurons as compact or compartmentalized, morphological features could not fully explain experimentally measured voltage distributions. This discrepancy may reflect limitations in precisely modeling the pattern of synaptic inputs evoked by the stimulus *in vivo*, or potential contributions from other molecular features. Future studies both *in silico* and *in vivo* will be needed to reveal how these mechanisms contribute to the diversity of cell-type specific voltage distributions.

### Voltage compartmentalization greatly expands the computational capacity of single neurons

Our modeling and experimental results demonstrate that cell-type specific electrical properties are not binary: neurons are not only isopotential nor wholly independent subcellular compartments but instead occupy a diverse continuum of subcellular electrical distributions. These modes of voltage distribution directly impact neuronal computation. For electrically compact cell types the electrical signals across each neuron are integrated simultaneously, producing voltage distributions and receptive fields that are uniform over all neurites and output synapses (Fig.6 and Extended Data Figure 12). This uniform distribution mandates all-to-all interactions amongst inputs, no matter where they are located on the cell’s surface, creating a single unit of computation at the level of membrane potential.

For electrically compartmentalized cells, electrical signals across each neuron can be integrated locally. In Dm9, CT1 and Melo13, this local integration produces distinct spatial receptive fields across neuronal arbors (Fig.6 and Extended Data Figure 15). These observations of local receptive fields could account for some of the discrepancies between connectomic predictions and direct measurements of receptive field size^61,62^. These non-uniform voltage distributions enable selective interactions amongst synaptic inputs within sub-cellular compartments, creating a substrate for parallel processing in single neurons. As output synapses are widely distributed in most cell types in the visual system, including across these sub-cellular compartments, these non-uniform voltage distributions can be translated into multiple different outputs (Extended Data Figure 16).

Our observations of a continuum of electrical properties, combined with previous work, suggest multiple mechanisms by which neurons might process and transmit information to downstream partners^14,22,24,25,30-32,35^. In compact neurons, the uniform changes in membrane potential across output synapses could result in each synapse relaying the same information to downstream targets. However, through localized differences in how membrane potential is coupled to calcium entry, compact neurons could also relay distinct information to different downstream targets. For compartmentalized neurons, different output synapses can sample from distinct voltage signals, meaning that either uniform or compartmentalized voltage to calcium transformations could be used to convey distinct information to downstream targets. Finally, while the regulated transformation of membrane potential to calcium signals can be used to threshold, compartmentalization allows a much more flexible platform for shaping signal transmission to multiple outputs. For example, the multiple receptive field structures seen in Dm9 and MeLo13 can only be achieved by voltage compartmentalization.

Voltage compartmentalization also raises the possibility that there are rich interactions between subcellular electrical compartments, allowing a single neuron to hierarchically integrate signals across different scales. By selectively compartmentalizing particular input combinations, a neuron can implement specific computational orders of operation. In the visual system, such a mechanism could enable a single neuron to perform sophisticated computations such as parallel feature extraction, multi-scale integration, and flexible normalization, computations that were previously only thought possible at the network level^63-66^. Future work can further explore the possibility of cross-compartment interactions that support sophisticated multi-input, multi-output computations within individual neurons^67^. Such fundamental insights into neuronal information processing will in turn inform designs of neuromorphic circuits and artificial neural networks^68-72^.

## Methods

### Animals

All flies used for imaging were raised on standard molasses food at 25°C on a 12/12h light-dark cycle. For sensor comparison, ASAP7y was inserted into the attP40 PhiC31 landing site by injection of a pJFRC7-20XUAS-ASAP7y plasmid (BestGene). For voltage imaging of dendrites, ASAP7y was put into SPARC cassettes with D (50%), I (10%) or S (1%) labeling densities and injected into the su(Hw)attP5 landing site using CRISPR (BestGene) as previously described^40^. Transformants were crossed to w-; S/CyO(Cre); Tm2/Tm6 to cut out the dsRed marker used in the CRISPR injection before being crossed with Gal4 driver lines.

For sensor comparison, we used the cell-type-specific driver GMR19F01-Gal4 to express GEVIs in Mi1, and R60H12-Gal4 to express GEVIs in TPN-II neurons. Female flies of the appropriate genotype were collected on day 2 and imaged on day 7 post eclosion.

Mi1>ASAP5: +/+; 20XUAS-ASAP5/+; + /GMR19F01-Gal4

Mi1>ASAP7y: +/+; 20XUAS-ASAP7y/+; + /GMR19F01-Gal4

TPN-II>ASAP5: +/+; 20XUAS-ASAP5/+; +/GMR60H12-Gal4

TPN-II>ASAP7y: +/+; 20XUAS-ASAP7y/+; +/GMR60H12-Gal4

For voltage imaging of neurites in visual neurons, all split-driver lines were crossed with nSyb-PhiC31; S/CyO; Tm2/Tm6B to include the nSyb-PhiC31 necessary for SPARC labeling^74-76^. We crossed the resulting nSyb-PhiC31-containing lines to SPARC-I-ASAP7y flies (generated as described above) to achieve sparse expression of ASAP7y in genetically defined cell types. Female flies of the appropriate genotypes were collected on day 2 and imaged on days 3-14 post eclosion, depending on the genotype.

LC4: nSyb-PhiC31/+; 20XUAS-SPARC-I-ASAP7y/GMR47H03-AD; + /GMR72E01-DBD

LC11: nSyb-PhiC31/+; 20XUAS-SPARC-I-ASAP7y/ R22H02 -AD; +/ R20G06 – DBD

LPLC2: nSyb-PhiC31/+; 20XUAS-SPARC-I-ASAP7y/GMR19G02-AD; + /GMR12E04-DBD

LC14b: nSyb-PhiC31/+; 20XUAS-SPARC-I-ASAP7y/ R44A02-AD; + / R50B11-DBD

Dm9: nSyb-PhiC31/+; 20XUAS-SPARC-I-ASAP7y/GMR10E07-AD; + /VT017422-DBD

Y3: nSyb-PhiC31/+; 20XUAS-SPARC-I-ASAP7y/R94E06 - AD; +/ VT026017- DBD

MeLo13: nSyb-PhiC31/+; 20XUAS-SPARC-I-ASAP7y/VT048356-AD; + /GMR16E12-DBD

CT1: nSyb-PhiC31/+; P[20XUAS-ASAP7y]attP40/R65E11-AD; P[20XUAS-ASAP7y]VK00005/R20C09-DBD.

For rat cortical neuronal cultures, Sprague Dawley rat (001 Charles River) embryos at embryonic day 18 were obtained with the approval of the Stanford Administrative Panel on Laboratory Animal Care (APLAC). Male and female rats were used interchangeably. Other mouse procedures are detailed in separate sections below.

### Scanless two-photon voltage imaging

#### Microscope

All scanless two-photon voltage imaging was performed using temporally focused, holographic spots that had a 12-µm lateral extent and approximately a 14-µm axial extent as described previously^48^. Excitation was provided by a tunable femtosecond source (Coherent Discovery, 1 W, 80 MHz, 100 fs) tuned to 1030 nm, with power externally modulated using a combination of half-wave plate and polarizing beamsplitter cube (Thorlabs, WPHSM05-980 and PBS253 respectively). The initial 3-mm diameter beam was expanded 2× using a telescope formed of a 100-mm focal length lens (Thorlabs, LC1093-C) and a second lens of 200-mm focal length (Thorlabs LA1979-C-ML) to underfill a spatial light modulator (SLM, Hamamatsu, LCOS X10468, 600 × 800 pixels, 20-μm pitch). Phase masks optimized to obtain a single holographic spot of diameter 12 µm at the center of the field of view were generated with an iterative Gerchberg-Saxton algorithm. Two-photon excited fluorescence was collected using a simple widefield detection axis comprised of the same objective lens and a tube lens (Thorlabs, TTL200-A) and a scientific complementary metal-oxide semiconductor (sCMOS) camera (Photometrics, Kinetix) located in a conjugate image plane. The camera was controlled with Micro-manager 2.0. Excitation and fluorescence were separated spectrally using a combination of dichroics (Chroma, ZT405/488/561/640rpc and Chroma T680spxr), band-pass filter (Semrock, FF03-525/50-25) and short-pass filter (Semrock, FF01-750sp). Acquisition rates for all datasets are specified throughout the manuscript.

#### Sample preparation

##### Organotypic hippocampal slices

All experimental procedures were conducted in accordance with guidelines from the European Union and institutional guidelines on the care and use of laboratory animals (Council Directive 2010/63/EU of the European Union). Organotypic hippocampal slices were prepared from mice (Janvier Labs, C57Bl6J) at postnatal day 9. Hippocampi were sliced with a tissue Chopper (McIlwain type 10180, Ted Pella) into 300 µm thick sections in a cold dissecting medium consisting of GBSS (Sigma, G9779) supplemented with 25 mM D-glucose, 10 mM HEPES, 1 mM Na-Pyruvate, 0.5 mM α-tocopherol, 20 nM ascorbic acid, and 0.4% penicillin/streptomycin (5000 U/mL). After 45 min of incubation at 4°C in the dissecting medium, slices were placed onto a porous membrane (Millipore, Millicell CM PICM03050) and cultured at 37 °C, 5% CO_2_ in a medium consisting of 50% Opti-MEM (Fisher 15392402), 25% heat-inactivated horse serum (Fisher 10368902), 24% HBSS, and 1% penicillin/streptomycin (5000 U/mL). This medium was supplemented with 25 mM D-glucose, 1 mM Na-Pyruvate, 20 nM ascorbic acid, and 0.5 mM α-tocopherol. After three days in vitro (DIV), the medium was replaced with 82% neurobasal-A (Fisher 11570426), 15% heat-inactivated horse serum (Fisher 11570426), 2% B27 supplement (Fisher, 11530536), 1% penicillin/streptomycin (5000 U/mL), 0.8 mM L-glutamine, 0.8 mM Na-Pyruvate, 10 nM ascorbic acid and 0.5 mM α-tocopherol. This medium was removed and replaced once every 2–3 days. Slices were transduced with AAV9-hSyn-ASAP7y at DIV 3 by bulk application of 1 µL of virus per slice (Final titer: 4.4 × 10^11^ vg/mL). Experiments were performed between DIV 10 and 17.

#### Electrophysiology

An upright microscope (Scientifica, SliceScope) was equipped with a far-red LED (Thorlabs, M660L4), oblique condenser, microscope objective (Nikon, CFI APO NIR, 40×/0.8-NA), tube lens (Thorlabs, TTL200-A), and an sCMOS camera (Photometrics, Kinetix) to collect light transmitted through the sample. Patch-clamp recordings were performed using an amplifier (Molecular Devices, Multiclamp 700B), a digitizer (Molecular Devices, Digidata 1550B) at a sampling rate of 10 kHz and controlled using pCLAMP11 (Molecular Devices). Cells were continuously perfused with artificial cerebrospinal fluid (ACSF) comprised of 125 mM NaCl, 2.5 mM KCl, 1.5 mM CaCl_2_, 1 mM MgCl_2_, 26 mM NaHCO_3_, 0.3 mM ascorbic acid, 25 mM D-glucose, 1.25 mM NaH_2_PO_4_. Continuous aeration of the recording solution with 95% O_2_ and 5% CO_2_ resulted in a final pH of 7.4 (as measured). For experiments on organotypic hippocampal slices, the ACSF was supplemented with 1 µM of AP5 (abcam, ab120003) and NBQX (abcam, ab120046), except for the spontaneous activity recordings. Borosilicate pipettes (OD 1.5 mm, ID 0.86 mm, length 10 cm, fire-polished with filament, WPI) were pulled using a pipette puller (P1000, Sutter Instruments) to a tip resistance of 4–6 MΩ. Pipettes were filled with an intracellular solution consisting of 135 mM K-gluconate, 4 mM KCl, 4 mM Mg-ATP, 0.3 mM Na_2_-GTP, 10 mM Na_2_-phosphocreatine, and 10 mM HEPES (pH 7.35). All reported membrane potentials reported were Liquid Junction Potential (LJP) corrected by –15 mV (measured). Recordings were compensated for capacitance (Cm) and series resistance (Rs) to 70%. Only recordings with an access resistance below 30 MΩ were included in subsequent analysis.

#### Imaging

To assess the compatibility of ASAP7y with scanless two-photon voltage imaging (**Extended data figure 3**), CHO cells were patched and clamped at –70 mV, at 21–23 °C, 48–72 hours post-transfection. A 20-Hz train of ten 3-ms 100-mV steps was electrically induced, and fluorescence was simultaneously imaged at 993 fps for a duration of 500 ms with irradiance of 0.88 mW/µm^2^, corresponding to 100 mW/cell. To measure the F-V curve of ASAP7y under scanless two-photon illumination at 1030 nm (**Extended data figure 3**), ASAP7y-expressing CHO cells were patched and clamped at −70 mV, and 300 ms voltage steps were applied (ranging from –110 mV to +30 mV). Scanless two-photon illumination was synchronized (duration of 400 ms centered on the voltage steps, irradiance 0.88 mW/µm^2^ corresponding to 100 mW/cell) and fluorescence was imaged at 100 fps. The ability to record APs in neurons in organotypic slices was assessed by electrically triggering AP trains for 1 s at 10–100 Hz while fluorescence was recorded by scanless two-photon imaging at 500 fps with irradiance of 0.44–1.11 mW/µm^2^, corresponding to 50–125 mW/cell. Spontaneous activity of neurons in organotypic slices was simultaneously electrically monitored and optically recorded by scanless two-photon imaging at 500 fps for 30 s with irradiance 0.88 mW/µm^2^, corresponding to 100 mW/cell. Imaging data were analyzed as previously described^48^.

### *In vivo* patch-clamp electrophysiology

#### In utero electroporation of mice

All animal procedures were approved by the Columbia University Institutional Animal Care and Use Committee (IACUC Protocol #AC-AAAB3562) and were conducted in accordance with NIH guidelines. In utero electroporation was performed at embryonic day (E) 15.5–16.5 following previously published protocols^15,79^. Briefly, pregnant CD-1 mice (Charles River) were anesthetized with isoflurane (3% for induction, 2% during surgery). Prior to surgery, animals received a local subcutaneous injection of lidocaine (2 mg/kg) and carprofen (5 mg/kg) was administered in the drinking water beginning 24 h before the procedure. Postoperative analgesia was maintained by providing carprofen in the water for an additional 2 days. A midline laparotomy was performed to expose the uterine horns. Plasmid DNA (1.0 μg/μL) mixed with Fast Green (0.05% w/v) was injected into the left lateral ventricle of each embryo using a pulled glass micropipette. For sparse labeling of Baz1a-positive neurons, plasmid mixtures included EF1α-DIO-ASAP7y in combination with a Baz1a-Cre plasmid. Throughout the procedure, the uterine horns were kept moist with warm sterile DPBS. Electroporation was carried out using platinum disk electrodes (Nepa Gene CUY650P5) positioned laterally on the embryonic head, together with a second electrode placed over the dorsal telencephalon (Nepa Gene CUY7003L). Electrical pulses were delivered using a Nepa Gene NEPA21 electroporator. After electroporation, the uterine horns were placed back into the abdominal cavity, and the abdominal wall and skin were closed with sutures.

#### Cranial window implantation for *in vivo* imaging and electrophysiology in mice

Electroporated mice (4–8 weeks old, both sexes) were anesthetized with isoflurane (3% for induction, 2% during surgery). Prior to the procedure, animals received a local subcutaneous injection of lidocaine (2 mg/kg), followed by intraperitoneal administration of enrofloxacin (5 mg/kg), dexamethasone (0.6 mg/kg), and carprofen (5 mg/kg). Postoperative analgesia was maintained with carprofen (5 mg/kg, once daily for 2 days). A stainless steel headplate was affixed to the skull using cyanoacrylate adhesive and dental cement, positioned above the left primary somatosensory cortex (S1) (centered at −3 mm lateral and −1 mm anteroposterior from bregma). A rectangular craniotomy (1.5 × 3 mm) was opened at the same coordinates, and the dura mater was carefully removed across the entire window. To minimize motion artifacts and enable two-photon imaging during whole-cell patch-clamp recordings, the craniotomy was sealed with a custom coverslip (1 × 4 mm). The coverslip was positioned to leave approximately 0.5 mm of exposed brain at the posterior edge to allow pipette access for targeted patch-clamp recordings.

#### *In vivo* two-photon-guided whole-cell electrophysiology in mice and simultaneous two-photon voltage imaging

*In vivo* two-photon-guided whole-cell patch-clamp recordings were performed in 4–8-week-old mice under isoflurane anesthesia (1–1.5% during recordings), following established protocols^15,80^. S1 layer 2/3 Baz1a-positive pyramidal neurons were targeted based on their location and fluorescence in the two-photon imaging plane. These neurons were electroporated to express genetically encoded voltage indicators for simultaneous two-photon voltage imaging. Patch pipettes (7–9 MΩ), pulled from borosilicate glass capillaries (1.5 mm OD, 0.86 mm ID; Sutter Instruments) using a horizontal puller (Sutter P-97), were filled with an internal solution (in mM: 126 K-gluconate, 20 KCl, 2 MgCl₂, 10 HEPES, 10 phosphocreatine, 4 Mg-ATP, 0.3 Na-GTP, 0.02 Alexa Fluor 594; pH 7.31; 298 mOsm). A high positive pressure (∼300 mBar) was maintained during pipette advancement to prevent clogging, then reduced to 50–100 mBar once ∼100 μm below the pia to navigate through cortex under two-photon guidance. Upon contacting the target soma, pressure was lowered to 10–20 mBar to indent the membrane and facilitate giga-seal formation, and whole-cell access was established after breakthrough. Recordings were accepted only if access resistance stabilized between 50–100 MΩ. Neurons were voltage-clamped at −60 mV with series-resistance compensation, and signals were amplified (MultiClamp 700B, Molecular Devices), digitized at 10 kHz, and acquired using PrairieView 5.8 (Bruker) software synchronized to imaging. Two-photon imaging was performed on a custom-built microscope (modified Prairie/Bruker inverted design) equipped with a 25×/1.05-NA water-immersion objective (Olympus) and a Ti:Sapphire femtosecond laser (Chameleon Ultra II, Coherent; 80 MHz, tunable 680–1080 nm) tuned to 1000 nm (for ASAP7y). Emitted fluorescence passed through a 525/50-nm filter (green channel) or a 605/15-nm filter (red channel for Alexa Fluor 594) and was detected by gallium arsenide phosphide (GaAsP) photomultiplier tubes. Frame-scan time series (256×121 pixels) were acquired at ∼120-Hz via a resonant galvanometer scanner to capture membrane voltage dynamics. Neurons were held at −60 mV and stepped from −120 mV to +40 mV in 20-mV increments (five sweeps per cell) to assess the voltage indicator’s sensitivity.

#### Imaging processing and fluorescence quantification for *in vivo* in mice calibration

Two-photon imaging movies were rigidly motion-corrected using Suite2P, and fluorescence time series were extracted from manually drawn somatic membrane ROIs using ImageJ. Imaging timestamps from PrairieView XML metadata provided a time base for aligning fluorescence signals. Raw fluorescence intensities for each ROI were normalized to local background levels measured in adjacent neuropil (via a soma-sized background ROI) to correct for baseline fluorescence and scattering, and a low-pass filtered (fourth-order, 0.1 Hz) version of this background trace was subtracted from each ROI’s signal to remove slow global drift. The fluorescence of all pixels within each ROI was averaged, and prior to ΔF/F analysis, the resulting traces were smoothed using a Savitzky–Golay filter (2nd-order, 5-ms window, adjusted to nearest odd frame count based on acquisition rate) to reduce high-frequency noise and stabilize baseline estimation. All image analysis was performed with custom routines in Python, and no blood autofluorescence or other artifacts were observed. Inverted sensor signals were treated as fluorescence changes (ΔF). Simultaneously recorded voltage-clamp command traces were imported from the electrophysiology files and aligned to the imaging time base. For each voltage step and ROI, we quantified the fluorescence response using a standardized ΔF/F method: baseline fluorescence F₀ at –60 mV was calculated from a 500-ms pre-step window, and within the voltage step (excluding ±10 ms around onset and offset to avoid edge artifacts), the fractional fluorescence change ΔF/F (relative to F₀ at –60 mV) was computed, with sign convention adjusted for the inverse-polarity sensor. The response amplitude for each step was defined as the sign-aware peak ΔF/F within this guarded window (maximum for depolarizing steps, minimum for hyperpolarizing steps), and the corresponding peak time and raw fluorescence value were recorded.

### SLAP2 voltage imaging

C57BL/6J male mice (#000664 Jackson Laboratory) of 2 months of age were anesthetized and placed in a stereotactic mount. Following craniotomy and dura removal, viral injections were performed at two sites in the left visual cortex area V1 (−3.1 A/P, -3.1 M/L, 0.5 D/V and -3.9 A/P, -3.1 M/L, 0.5 D/V), using a Nanoject III injector and a beveled borosilicate micropipette. Sparse expression was achieved by injecting 200 nL of high-titer Cre-dependent indicator and low-titer Cre AAV mix: PHP.eB-hSyn-DIO-ASAP7y (1 × 10^12^ vg/mL) + PHP.eB-hSyn-FLEX-HaloTag-WPRE (1 × 10^12^ vg/mL) + PHP.eB-CamKII0.4.Cre.SV40 (1.2 × 10^8^ vg/mL). *In vivo* imaging was carried out after at least 4 weeks of expression and 5 days of habituation for head fixation. Mice were not anesthetized during imaging and were allowed to run freely on a wheel. The imaging system is previously described^47^ using a Leica 25×, 1.0 NA objective. Excitation was done at 1030 nm and post-objective power was 35 mW. All the ROIs were uniformly scanned at 2 kHz. Visual stimuli were full-field square-wave drifting gratings presented in 8 directions (0–315°, 45° steps), with a spatial frequency of 0.04 cycles/degree and a temporal frequency of 2 Hz. Each stimulus lasted 2 s and was followed by 1 s of a gray screen.

### Mesoscope Imaging

#### Surgery

Four-week-old mice were retro-orbitally injected with 1 × 10^11^ vg per mouse of AAV PHPeB-CaMKIIa-ASAP7y-Kv2.1, produced by the Stanford Gene Vector and Virus Core. Widefield imaging preparations were performed as described previously^81^. Briefly, mice were anesthetized with isoflurane, and the scalp was removed. The skull was cleaned, and a custom headplate and scope holder were affixed using dental cement. The exposed skull was then coated with a thin layer of cyanoacrylate glue followed by clear nail polish. For postoperative analgesia, mice received a single subcutaneous injection of sustained-release buprenorphine (0.1 mg/kg). Mice were allowed to recover for at least one week before the start of experiments. One day prior to recording, mice were anesthetized with isoflurane and two craniotomies were performed, one for the silicon probe and one for an electrical reference affixed with dental cement. The open craniotomy was covered with silicone rubber overnight.

#### Recordings

Animals were habituated to head fixation before experiments. Head fixed mice were recorded with a silicon probe placed in motor cortex and a reference electrode placed in the olfactory bulb. Fluorescence was recorded using a tandem lens microscope with an f/1.2 aperture and 510-nm illumination. Illumination power was 20 mW across the whole brain, with an irradiance of 200 μW/mm^2^.

#### Processing

To calculate correlation coefficients and coherence, 1-minute long intervals were sampled from 8-minute continuous recordings. Plots and reported numbers are the average and standard error of the mean of these samples. Video data, and data for Pearson correlation was bandpass filtered from 1 to 60Hz.

### Fiber optic imaging (TEMPO)

#### Viral vectors

For fiber-optic TEMPO studies we expressed GEVIs using AAV9-EF1α-DIO-ASAP7y (4 × 10^12^ GC/mL) or AAV2/PHP.eB- EF1α-DIO-ASAP3 (1.3 × 10^13^ GC/mL) and a reference fluorophore via AAV2/PHP.eB-CAG-mRuby2 (2 × 10^14^ GC/mL).

#### Pharmacology

For studies of anesthetized mice, we used ketamine (VEDCO #50989-996-06) and xylazine (AnaSed #59399-110-20). We dissolved ketamine-xylazine in phosphate buffered saline (PBS; 10 mg/mL ketamine; 1 mg/mL xylazine), yielding final dosages of 100 mg/kg and 10 mg/kg, respectively. We injected these drugs intraperitoneally with a 30-gauge needle at least 20 min before data acquisition.

#### Mouse surgery

Animal procedures were approved by the Stanford APLAC. We used ∼12-week-old male or female mice from the PV-Cre driver line (PV-IRES-Cre; Jackson Laboratory #008069). We housed mice in normal light cycle conditions, 2–5 per cage before surgery, and 1 per cage afterward. Mice were provided with food and water *ad libitum*. Animal surgery was conducted as previously described^50^. Briefly, mice underwent two surgical procedures under isoflurane anesthesia (1.5%–2% in O_2_). In the first procedure, we injected viruses to express the GEVI and the reference fluorophore. In the second procedure, performed about a week after viral injection, we implanted an optical fiber (400 µm core diameter, 0.5 NA, Thorlabs). Coordinates for virus injections in V1 were –3.5, 2.5, –1.2 (in mm from Bregma; AP, ML, DV). We injected 0.5 µL per mouse of a solution containing 5E9 genome copies (GC) of the GEVI virus and 1E9 GC of the reference virus. For all mice, we prepared the skull by manually removing the conjunctive tissue. We cleaned the skull by applying H_2_O_2_, followed by rinsing with Ringer’s solution (ThermoFisher #50980245). We drilled a 0.5-mm-diameter opening in the skull above V1 using 0.5-mm-diameter micro drill burr (Fine Science Tools #19007-05). We inserted the implant and sealed the gap between the skull and the implant using adhesive cured with ultraviolet light (Loctite 33105). We glued a custom-designed stainless steel head bar onto the mouse skull and secured the implant to the skull using blue-light cured resin (Flow-it ALC, Pentron). To mitigate post-operative pain, we administered carprofen (5 mg kg^−1^) about 30 min prior to the end of surgery. Mice recovered for >2 weeks before imaging experiments began.

#### Visual stimulus presentation

We placed an LCD monitor (85-cm diagonal) in the monocular visual field at a distance of 20 cm, contralateral to the fiber optic implant. We used custom software in Psychtoolbox (MathWorks) to display drifting gratings (sine waves of 1 Hz temporal frequency, 0.033 cycle per degree of spatial frequency and 100% contrast, oriented at 315° to horizontal). We placed a blue bandpass optical filter (#5084 Damson Violet, Rosco E-Colour) onto the video monitor to ensure its emissions were outside the range of wavelengths detected by the TEMPO apparatus. The drifting grating was presented for 1.5 s and was preceded and followed by a gray isoluminant screen. Excitatino wavelengths used were 488 nm for ASAP7y and 561 nm for mRuby2 as previously described^50^. Mice underwent a total of 50 trials, with the inter-trial intervals randomized between 2–5 s.

#### Data processing for fiber-optic TEMPO recordings

For all fiber optic TEMPO recordings we analyzed the data using a custom-written uSMAART analysis package following previous methods (https://github.com/sihaziza/uSMAART_public) that runs in MATLAB (Mathworks)^50^. To compute cross-frequency coupling between two oscillations of different frequencies, we first bandpass-filtered the raw signal at the low-frequency bandwidth of interest. From this filtered version of the trace, we estimated the analytic signal using the Hilbert transform and computed the phase. Next, we estimated the wavelet spectrogram of the raw signal and computed the average across the high-frequency bandwidth of interest. Finally, we identified every phase reset of the low-frequency oscillation and aligned the high-frequency signal accordingly.

### Simulating electrotonic properties of neuronal reconstructions using NEURON

We simulated the electrotonic properties of all neurons in the connectome of the visual system of a male Drosophila^4^. We used the neuprint (https://connectome-neuprint.github.io/neuprint-python/docs/skeleton.html) Python client to fetch skeletons in SWC format. We scaled each skeleton with a geometric factor of 0.008 µm, as neuprint uses a coordinate system based on the grid of EM voxels (each voxel is an 8 nm cube). The SWC file was used as input to the NEURON (https://www.neuron.yale.edu/neuron/) software in Python, with electrical parameters close to previously estimated parameters from electrophysiological recordings^24,77^, to build NEURON models. If not otherwise specified, the default passive parameters were: specific membrane conductance G_pas_ = 0.1 × 10^-3^ S/cm^2^, specific membrane capacitance C_m_ = 1.0 µF/cm^2^, and intracellular resistivity R_a_ = 200 Ω· cm. The resting membrane potential was –65 mV. For each neuron, we used the 5 sections that had the highest density of input synapses from the most highly connected upstream neuron as input sections. We performed 5 simulations for each neuron, and for each simulation run, we chose one input section and injected 50 pA of current for 10 ms at the beginning of the simulation. We simulated each neuron for 250 ms in 0.25 ms steps. We then extracted the voltage trace over each section of the neuron to compute its integrated voltage change over time.

#### Selection of simulation parameters

We chose the biophysical parameters (R_a_, G_pas_ and C_m_) and stimulation parameters (duration and amplitude of current injections) based on previously estimated biophysical parameters^24,77^, cable theory, and empirical tests with Y3 neurons. As passive cable theory uses a linear equation, we reasoned that the duration and the amplitude don’t affect the relative voltage decay between two regions on a neuron. Indeed, when we varied the duration of the current injection from 5 ms to 40 ms, the relative voltage decay between two regions (medulla and lobula) of Y3 neurons remained unchanged (**Extended Data Figure 7**). Cable theory also suggests that the voltage decay is controlled by a space constant 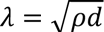 where *d* is the diameter of the cable and *ρ* = 1/(4 × *R_a_* × *G_pas_*) is a constant determined by R_a_ and G_pas_ but not C_m_. Indeed, we observed that by varying C_m_, the relative voltage decay between two regions (medulla and lobula) of Y3 neurons remained unchanged (**Extended Data Figure 7**). Indeed, when we varied R_a_ or G_pas_ while keeping their product the same, the relative voltage decay between two regions (medulla and lobula) of Y3 neurons also remained unchanged (**Extended Data Figure 7**). Taken together, these results suggest that the main biophysical parameter that affect voltage decay in a passive neuron model is *ρ* = 1/(4 × *R_a_* × *G_pas_*).

#### Simulating all neuron types in the optic lobe

We simulated all neuron types in the optic lobe. To capture high-quality reconstructions, we selected 5 neurons with highest number of reconstructed coordinates in the downloaded SWC files for each neuron type. If a neuron type has less than 5 neurons, we selected all of them in the simulations. The simulations were performed as described above.

#### Quantification of geometrical features

For each simulation of a neuron, we quantified the geometrical features based on the neurite connecting the injection site and the output segment, defined as the segment with output synapse with the lowest voltage signal. For contralaterally projecting neurons, which were incompletely reconstructed, we used the neurite connecting the injection site and the segment with the lowest voltage signals in the reconstructed neuron. The path length *L* was computed along the neurite. To extract a representative radius of the neurite, we computed the weighted average of the radii *r*_eq_ of all segments according to the following formula

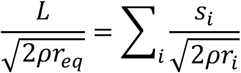

where *s*_i_, *r*_i_ is the length and radius of segment *i* and *ρ* is defined above. The intuition of this formula is that the space-constant-normalized path length is the sum of all space-constant-normalized segment lengths. The number of branches was counted along the neurite, where a branch from a NEURON section was defined as a child section that has more than 1 child and is not the next section along the neurite. We excluded the neurons that have a path length < 10 μm or that have no branch points. In total, we analyzed 717 neuron types.

#### Simulating neuron types measured experimentally

We simulated measured neurons LC4, LC11, LC14b, LPLC2, Y3, Dm9 and MeLo13 with more skeletons and across a range of biophysical parameters as described above. Here we omitted CT1 as there is only 1 such neuron in one optic lobe. To capture high-quality reconstructions, we quantified the total number of x-, y-, and z-coordinates in the SWC file associated with each neuron’s skeleton. Neurons within each cell type were then clustered into two groups using a Gaussian mixture model (GMM) implemented with the GaussianMixture class from *scikit-learn* in Python. The GMM was fit to the distribution of coordinate counts, and each neuron was assigned to the component for which it had the highest posterior probability. For downstream simulations, we selected the 20 neurons with the highest coordinate counts within the high-quality cluster, corresponding to the most densely reconstructed neurons. In cases where fewer than 20 neurons were available in the high-quality cluster, all neurons assigned to the high-quality cluster were used. We simulated neurons across several different biophysical parameters, choosing R_a_ to be 100, 150, or 225 Ω· cm, and G_pas_ to be 0.05 × 10^-3^ or 0.2 × 10^-3^ S/cm^2^ such that *ρ* = 1/(4 × *R_a_* × *G_pas_*) varied from 5.55 to 50.0 cm, close to one order-of-magnitude. Simulations of each neuron were performed as described above.

### *In vivo* voltage imaging in flies with a random access two-photon microscope

All flies were imaged with an acousto-optic deflector-based random-access microscope (AODscope, Karthala) and a ×25 1.1-NA objective (Nikon N25X-APO-MP CFI LWD). The AOD scope is equipped with a pulse picker that transmits laser pulses with a duty cycle of 600 ns out of 3000 ns (corresponding to 20% transmission). Visual stimuli were generated as previously described using FlyStim^78^. Briefly, the output of the blue LED from one projector (Lightcrafter 4500, Texas Instruments) was filtered through a 482/18-nm filter and projected onto a screen at 120 Hz. A photodiode (Thorlabs, SM05PD1A) was used to align the timing of the stimulus to image acquisition with millisecond precision.

### *In vivo* sensor comparisons in Mi1

For sensor comparison in Mi1, flies were imaged at 1030 nm (24 mW) or 940 nm (16 mW) without pulse-picking. For each neuron, 5 regions of interest (ROIs) measuring 5 × 15 × 20 μm were positioned at neurite of Mi1 neurons in the M10 layer. An ROI was placed outside of the fly to record the photons emitted by the projector for visual stimuli. Samples were scanned at 1041 Hz. The emission light was collected through a 525/50 nm filter. Visual stimuli were full-field 8-ms contrast increments or decrements (100% Michelson contrast, background=0.2, dark flash=0.0, bright flash=0.4 in FlyStim, 50 presentations each, in random order), interleaved in time by 500 ms presentations of a mean gray background. The fluorescence traces after the background subtraction were IIR-notch filtered at 120 Hz to remove residual bleed-through signals from the projector. The peak response (peak ΔF/F) or peak SNR was calculated for each trial instead of the stimulus-triggered average (STA). The baseline brightness F_0_ was the average photon count of all pre-stimulus windows across the recording. SNR was calculated empirically, and the noise was estimated as the standard deviation of the fluorescence signals in a 100 ms window before the onset of a stimulus.

### *In vivo* sensor comparisons in TPN-II

For sensor comparison in TPN-II, flies were imaged at 1030 nm (33 mW) or 940 nm (27 mW) without pulse-picking. For each neuron, 6 regions of interest (ROIs) measuring 4 × 12 × 16 μm were positioned at the axonal terminal of an individual neuron. Samples were scanned at 1041 Hz. The emission light was collected through a 525/50 nm filter. We detected action potentials based on z-scored Gaussian smoothed (σ=1) ΔF/F with a threshold of 2. Then we quantified the peak amplitude of detected APs with the original ΔF/F traces. The baseline brightness F_0_ was the median photon count across the recording. SNR was calculated empirically, and the noise was estimated as the standard deviation of the 50-Hz high-pass-filtered fluorescence signals after removing APs.

### *In vivo* multi-site voltage imaging

For voltage imaging along neurites, flies were imaged at 1030 nm with 13-21 mW power at 1041 Hz with 20% transmission as described above. For each neuron, regions of interest (ROIs) measuring 2 × 6 × 8 μm were positioned along the neurites of an individual neuron. First, the receptive field center (RFC) of each neuron was determined by a two-stage mapping process (see below). All subsequent visual stimuli were then centered on the RFC. Following stimulus presentations, a high-resolution Z-stack (0.58 μm × 0.58 μm × 1 μm or 0.58 μm × 0.58 μm × 2 μm) was taken to capture neuronal morphology.

#### Receptive Field Mapping

To obtain an initial estimate of each neuron’s receptive field, we presented a dark spot (r = 15°, intensity = 0.0) for 200 ms on a uniform grey background (intensity = 0.2) across a coarse spatial grid spanning φ = –30° to 20° and θ = –50° to 50°, in 10° steps, with two repetitions per location. Each presentation was separated by an 800-ms inter-stimulus-interval. For each location, the peak response was quantified as the mean ΔF/F within 150 ms after stimulus onset across both presentations. These responses were fit by a 2D Gaussian to estimate the location of the RFC. In the second stage of receptive field mapping, a small, dark spot (r = 4°, intensity = 0.0) was presented for 100 ms on a uniform grey background (intensity = 0.2) across a 24°-by-24° grid centered on the coarse receptive field estimate (φ = –12° to 12°; θ = –12° to 12°, 4° steps, two repetitions per location). Each presentation was separated by an 800-ms inter-stimulus-interval. As in the coarse mapping stage, the mean ΔF/F in the 150 ms following stimulus onset was measured for each location, and a 2D Gaussian was fit to these responses to estimate the neuron’s RFC.

#### Visual Stimuli Presentation

Following receptive field mapping, all visual stimuli were presented centered on the RFC of each neuron. All stimuli (intensity = 0.01) were presented against a uniform grey background (intensity = 0.2). First, a small, dark dot (r = 4°,) was presented for varying durations (8, 16, 32, and 64 ms), with conditions randomly interleaved. Each stimulus was presented 10 times, with a 500-ms inter-trial interval. Next, dark dots of varying radii (r = 2.5°, 5°, 10°, and 20°) were presented for 80 ms, with conditions randomly interleaved, an inter-trial interval of 800 ms, and 10 stimulus presentations per condition. Lastly, a series of dark annuli of varying radii (r_inner_= 2.5°, r_outer_ = 5°; r_inner_= 5°, r_outer_ = 10°; r_inner_= 10°, r_outer_ = 20°; r_inner_= 20°, r_outer_ = 40°) were presented for 80 ms, with conditions randomly interleaved, an inter-trial interval of 800 ms, and 10 stimulus presentations per condition.

#### Data pre-processing

Point-scan voltage imaging data were processed using custom Python code. For analyses at the single-ROI level, the raw photon count traces from individual ROIs were analyzed separately. For region-based analyses, raw photon-count traces from ROIs assigned to the same region were grouped by summing their photon counts. To standardize sampling across recordings, fluorescence traces were linearly interpolated and resampled to 1-kHz. Next, a 120-Hz IIR notch filter was applied to remove projector-related artifacts, and a 200-Hz low-pass Butterworth filter was applied to attenuate high-frequency multiplier noise. The filtered traces were then smoothed with a Gaussian kernel (σ = 5) to reduce point-to-point fluctuations while preserving response dynamics. Lastly, the photodiode synchronization signal was used to segment the fluorescence traces into individual stimulus-aligned trials. For each trial, fluorescence changes were converted to ΔF/F according to 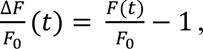 where *F_0_* is the mean fluorescence during the pre-stimulus baseline (time < 0).

#### Calculation of integrated signal, EUI, and slope

For each cell, trial wise ΔF/F traces were extracted from each region. To quantify the response magnitude, the integrated signal was computed for each trial. For a given trial, integration was performed over a fixed post-stimulus time window extending from stimulus onset (t = 0) to a cell-type specific end time derived from cell-type specific response dynamics (Dm9, 100 ms; MeLo13, 300 ms; Y3, 150 ms; LC14b, 200 ms, LC11, 400 ms; LPLC2, 200 ms). Only ΔF/F values that reflect depolarization were included in the integral. For each trial, we compared the integrated voltage responses measured across regions in the Electrical Uniformity Index (EUI). The EUI was computed as: 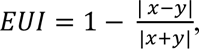 where *x* and *y* denote the trial-wise integrated signal measured in the two regions. For each cell, trial-wise integrated signals were fit using constrained linear regression of the form *y* = *mx* (intercept fixed at the origin). Slopes were estimated using ordinary least squares, and 95% confidence intervals were computed from the residual variance of the constrained model.

#### Comparing measured and modeled integrated signals

For each imaged cell type, we selected the 20 highest quality reconstructions (see **Simulating electrotonic properties of neuronal reconstructions using NEURON**) and defined morphological regions that corresponded to our experimental ROI placement. For each simulation run, we computed the average integrated signal within each region and then averaged these regional values across the five simulations of each neuron to compute the EUI for each neuron under each biophysical parameter set. Simulation results were grouped by biophysical parameters, and kernel density estimation (KDE) was used to visualize the distribution of ECIs across neuronal reconstructions.

#### Spatial tuning analysis

For analysis of spatial tuning across individual ROIs, trial-averaged ΔF/F traces were computed for each (θ, φ) condition from fine receptive field mapping (φ = –12° to 12°; θ = –12° to 12°, 4° steps, two trials per location), and response amplitudes within the 150-ms post stimulus onset window were used to generate ROI-specific tuning maps. A 2D gaussian was fit to these responses to estimate the ROI-specific RFC. To quantify the spatial offset between receptive field maps derived from different ROIs within the same neuron, we computed the 2D cross-correlation for each pair of receptive field maps. The location of the peak of the correlation matrix corresponds to the spatial shift that best aligns the two maps. The Euclidean norm of this shift was multiplied by the angular resolution of the mapping grid (4°) to yield the receptive-field shift in degrees of visual space. To account for the broad receptive field of CT1, the angular resolution of the mapping grid was 10°, and the dot size was 10°.

## Data Availability Statement

All source data will be made available from the corresponding authors upon request.

## Acknowledgements

We thank the Stanford Gene Vector and Virus Core for viral vector packaging, Lan Xiang Liu (Stanford University) for animal care. This work was supported by R01 EY022638, P30EY026877, The Chan Zuckerburg Biohub, San Francisco, a Stanford Bio-X Seed Grant to T.R.C, UM1MH136462 and 1RM1NS132981 to M.Z.L, L.J. was supported by the Stanford Graduate Fellowship and the National Defense Science and Engineering Fellowship, Y.A.H. was supported by the Stanford BioX Bowes Fellowship. B.B.A. and R.Y. were supported by the NINDS (RM1NS132981) and NEI (R01EY035248). C.C. was supported by a NIH Cellular and Molecular Training Grant (NIGMS, grant number 5T32GM007276).

## Author Contributions

Y.A.H. engineered and characterized ASAP7y, performed the fly experiments, performed simulations, analyzed data, prepared figures, and co-wrote the manuscript. L.L.J built flies, performed the fly experiments, contributed to simulations, analyzed data, prepared figures, and co-wrote the manuscript. S.L. characterized ASAP7y, made ASAP7y viruses and tested ASAP7y and variants in mice. M.N.D, Z.L and C.C engineered ASAP7y. M.Z. built flies, and imaged CT1 neurons in flies. S.H. performed TEMPO imaging. B.B.A. performed whole-cell patch-clamp *in vivo*. I.B. and R.R.S. performed scanless two-photon imaging. Y.A.H., M.S., A.N. performed SLAP2 two-photon imaging. A.D.W., J.K. and Y.W. performed transcranial mesoscopy with LFP. K.P. designed and constructed SLAP2 microscope. K.P., V.E., K.D., R.Y., J.D. and M.J.S. provided supervision, advised on experimental design, and assisted with data interpretation. M.Z.L. and T.R.C. conceived of the project, provided supervision, advised on experimental design, assisted with data interpretation, prepared figures, and co-wrote the manuscript.

