## Supplemental materials for "Ultrasensitive voltage imaging reveals distinct electrical microdomains in neurons"

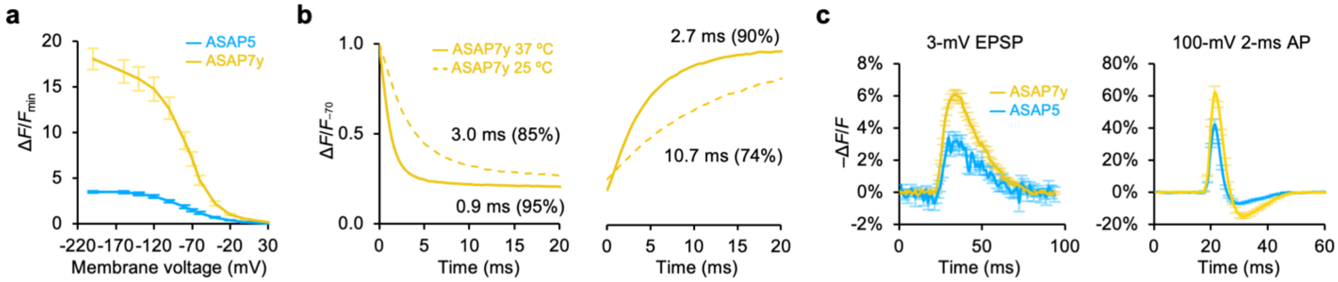


**Extended Data Figure 1. Additional characterization of ASAP7y *in vitro*.** **a**, Normalized fluorescence responses of GEVIs to their minimum fluorescence. *n* = 5 (ASAP5), *n* = 7 (ASAP7y) cells. Error bar: standard error of the mean. **b**, Activation dynamics (–70 mV to +30 mV) and deactivation dynamics (+30 mV to –70 mV) of ASAP7y in HEK293 cells. *n*=8 (25 °C) and *n* = 6 (37 °C). **c**, Fluorescence responses of GEVIs to commanded 3-mV EPSPs and 100-mV action potentials in HEK293 cells at 37 °C. *n* = 6 (ASAP5), *n* = 7 (ASAP7y) cells. Error bar: standard error of the mean.

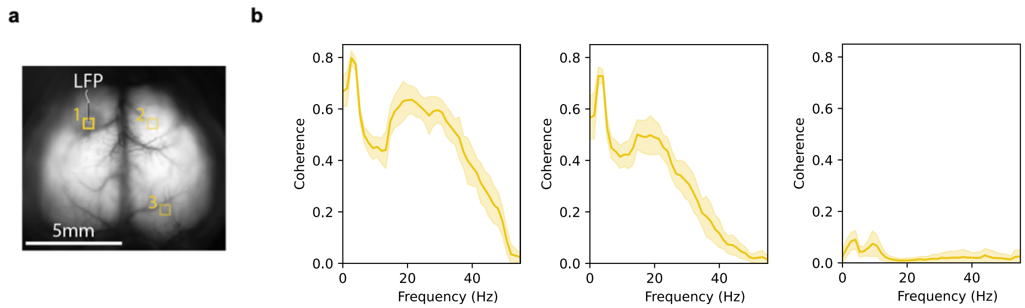


**Extended Data Figure 2.** Mesoscale voltage imaging with ASAP7y. **a,** Regions of interest (ROIs) 1, 2, and 3 from Fig. 1h. **b,** Coherence plots of correlation coefficients between LFP and ASAP7y signals from ROI 1, 2, or 3. Line and shaded area are mean ± SEM of eight 1-minute epochs in a continuous 8-minute recording. **c,** Fluorescence within ROI1 over time with constant illumination. Data are from one of two mice showing similar results.

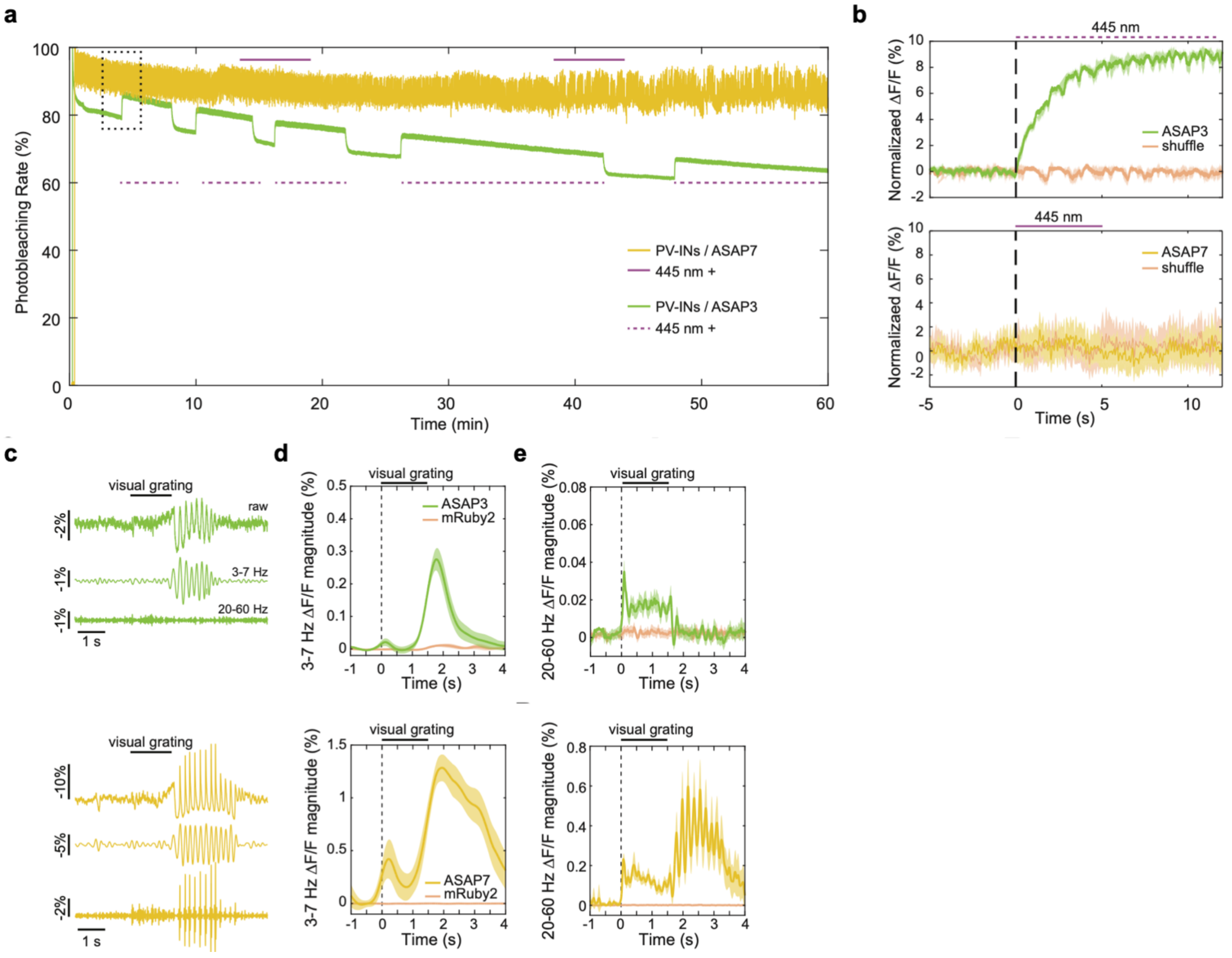


**Extended Data Figure 3. ASAP7y improves the detection sensitivity of TEMPO for cell-type specific high-frequency brain oscillations by ~10-fold. a,** Plot of photobleaching rate in a 1h-long recording from 2 mice expressing either ASAP3 or ASAP7y in PV interneurons. During the recording, bouts of CW 445 nm laser (~10 μW/mm$^2$ intensity) was delivered to the same brain location to estimate fluorescence recovery, as previously described for an earlier ASAP variants in Xu et al 2018 [PMID: 29866136]. After 1h-long illumination, we observed that ASAP7y exhibits minimal (~10%) photobleaching (compared to ~40% for ASAP3) and that ASAP3, but not ASAP7y, signal undergoes reversible photobleaching. Note that the ASAP7 trace demonstrates larger fluctuations due to the larger dynamic range of ASAP7y (~10% ΔF/F) as compared to ASAP3; these fluctuations are primarily gamma oscillations as further analyzed in **Fig. 1i**. The dashed black box indicates the period over which we quantified the metrics in (**Fig. 1i, j**). **b,** Plots quantifying the photoactivation effect for ASAP3 (top) and ASAP7y (bottom). While a 445 nm light exposure had no effect on ASAP7 fluorescence signal, it improved the ASAP3 baseline fluorescence $F_0$ by nearly 10%. Shading: C.I.95. **c,** ASAP7y (yellow) reports visually evoked gamma oscillations in PV interneurons of brain area V1 of awake mice with ~10-fold better SNR than ASAP3 (green). For each sensor, example traces of voltage dynamics of the raw (top), 3-7 Hz filtered (middle) and 20-60 Hz filtered (bottom) signals during a single trial of visual stimulation are shown. **d,** Top: Mean time-dependent fluorescence signal magnitudes in the 3-7 Hz (**c**) frequency bands for ASAP7y and the reference fluor mRuby2, computed using wavelet transforms and average over n=50 different trials. Shading: C.I.95. Bottom: Same as top but for the ASAP3-labeled PV interneurons. **e,** same as (**d**) but for gamma (20–60 Hz) frequency bands. Overall, combining ASAP7y and TEMPO improved the detectability of high-frequency gamma oscillations in awake mice by ~8 fold (visually evoked gamma magnitude in ΔF/F (%) of 0.016 ± 0.001 versus 0.124 ± 0.006 for ASAP3 and ASAP7, respectively, mean ± s.e.m. over n=50 trials each, rank sum test $p<0.0001$). Shading: C.I.95.

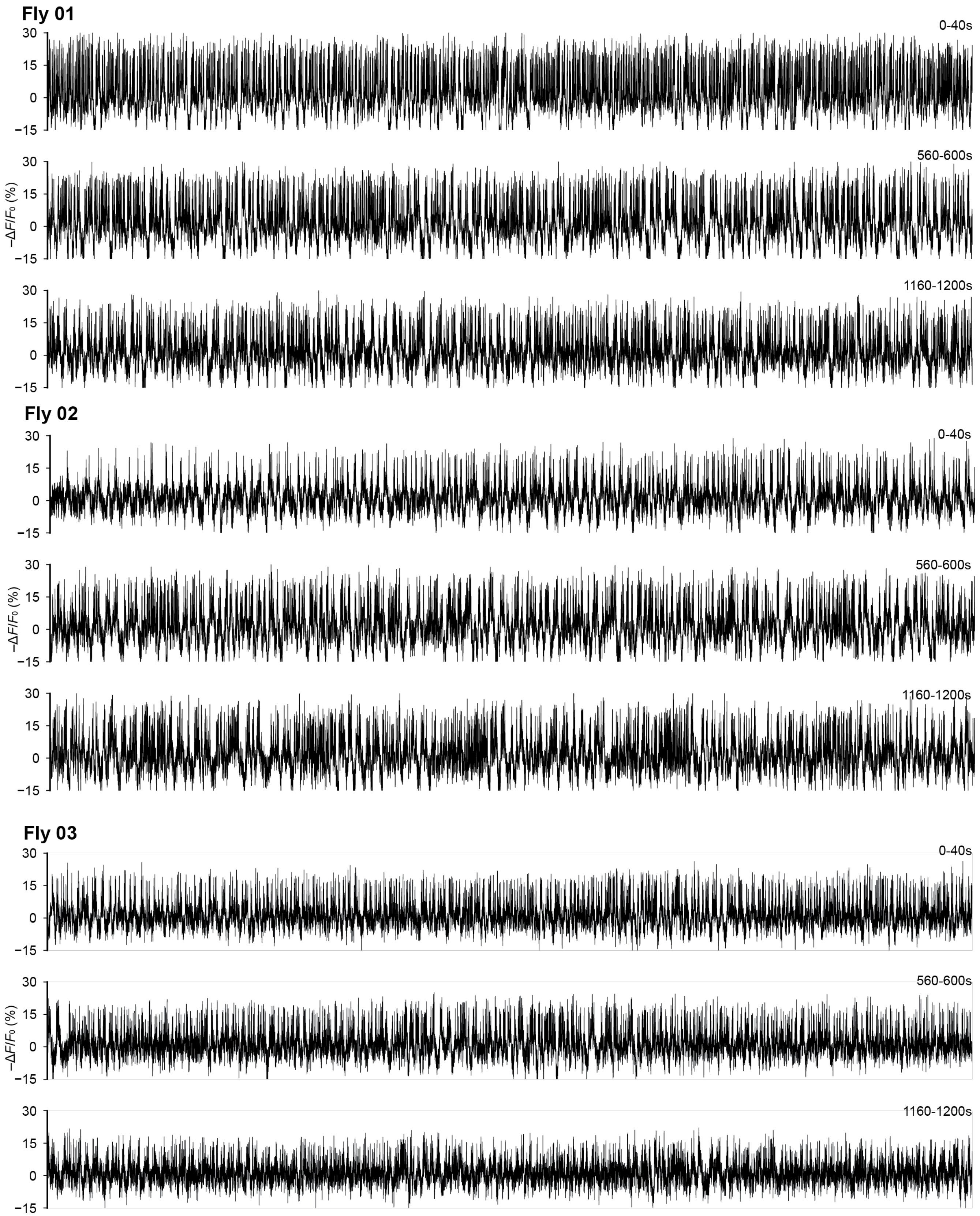


**Extended Data Figure 4. Extended Recordings of TPN-II neurons.** Fluorescence signals (–ΔF/F) of ASAP7y-expressing TPN-II neurons imaged continuously for 20 min at 1030 nm excitation. The first, middle and the last 40 seconds of the recording are shown, for 3 flies.

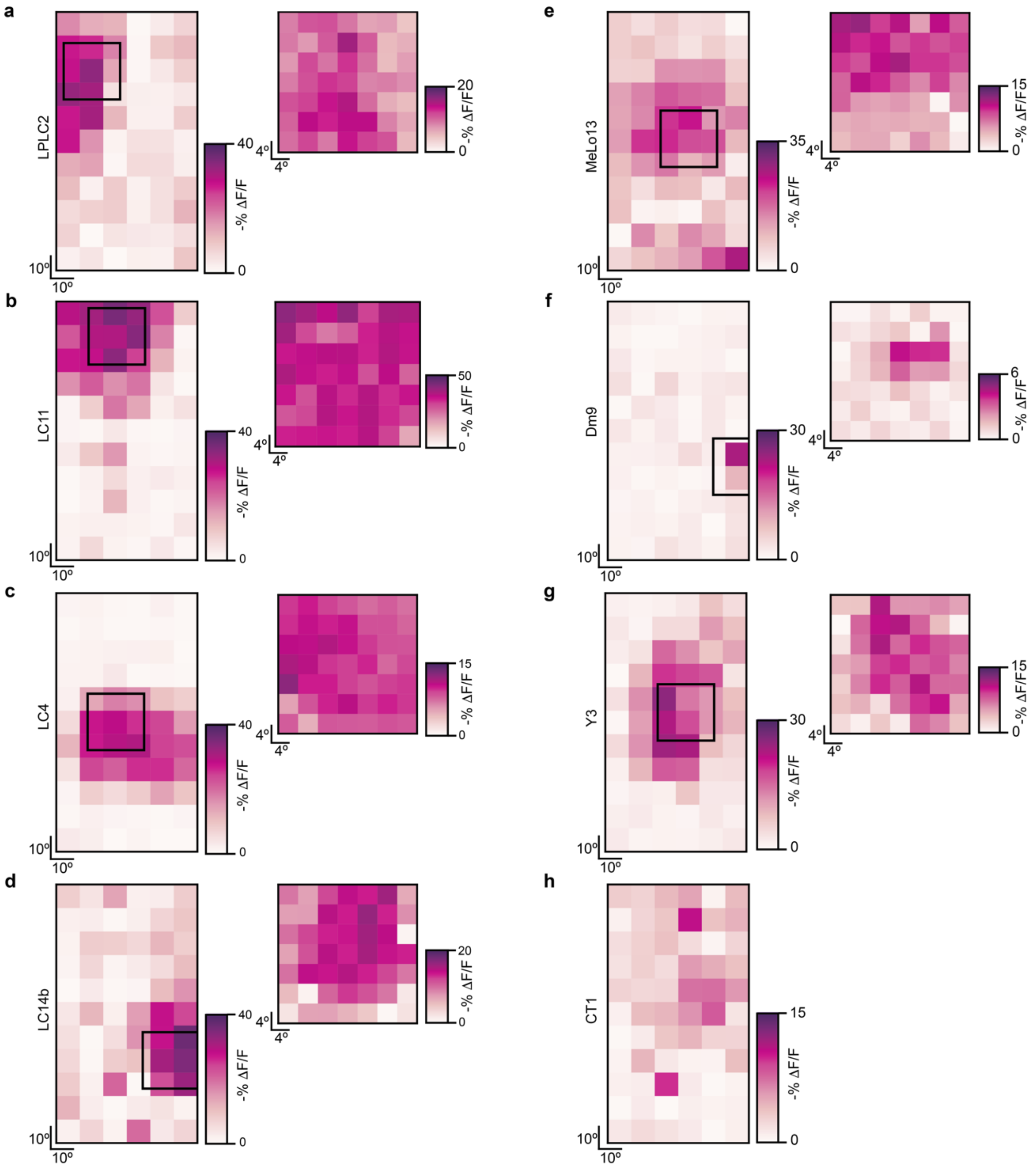


**Extended Data Figure 5. Receptive Field Mapping. a-h**, Receptive field mapping for cells presented in **Fig. 3** and **Extended Data Figure 6**. Left: responses from the first round of receptive field mapping. The grid for the second round of receptive field mapping is denoted by the black square. Right: responses from the second round of receptive field mapping.

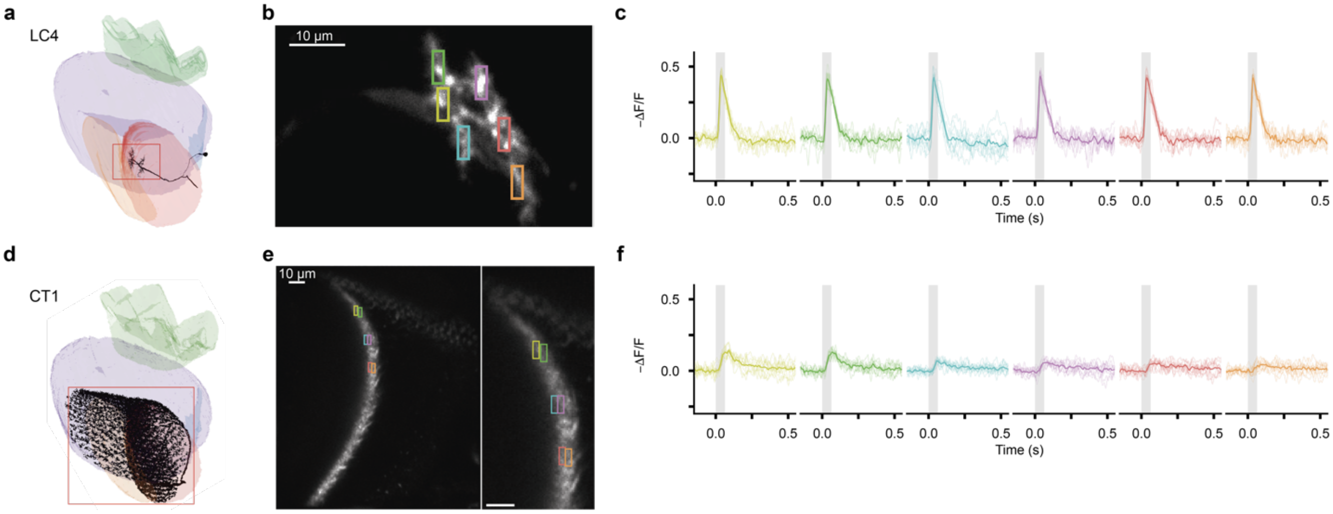


**Extended Data Figure 6. Single neuron responses in LC4 and CT1. a**, Reconstruction of a single LC4 neuron in the optic lobe. The red box indicates the imaging window. **b**, A single imaged LC4 neuron, with 2 × 6 *um* regions of interest (ROIs) overlaid. **c**, − ΔF/F responses from the ROIs shown in (**b**) over 10 stimulus trials, in response to a small spot stimulus (r = 4º, duration = 64 ms) presented at the neuron's receptive field center on an otherwise uniform grey background. Individual trials are shown as thin lines; the mean response over 10 trials is shown in bold; grey shading denotes the stimulus presentation window. **d-f**, same as (**a-c**) for CT1.

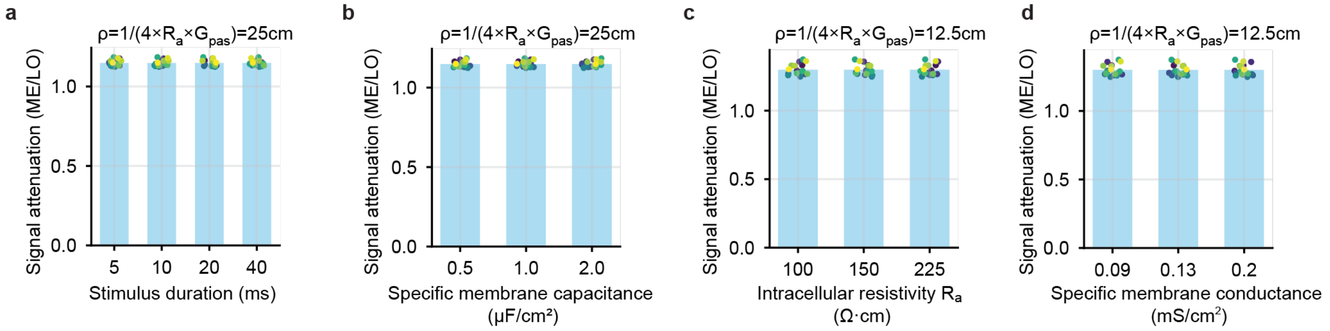


**Extended Data Figure 7. Selection of biophysical parameters for simulation. a**, Effect of duration of current injection on voltage decay: across 19 Y3 neurons simulated, the voltage decay measured as a ratio of the voltage in the medulla region and the lobula region is the same across 5, 10, 20, 40 ms of current injection. Each dot is an individual neuron. This is consistent with the linearity of passive cable theory. **b**, Effect of specific membrane capacitance on voltage decay: across 19 Y3 neurons simulated, the voltage decay measured as a ratio of the voltage in the medulla region and the lobula region is the same across 0.5, 1.0, and 2.0 μF/cm$^2$. This is consistent with passive cable theory. **c**, Effect of intracellular resistivity $R_a$ on voltage decay when the product of $R_a$ and membrane conductance $G_{pas}$ is fixed: across 19 Y3 neurons simulated, the voltage decay measured as a ratio of the voltage in the medulla region and the lobula region is the same across 100, 150, and 225 Ω·cm. **d**, Effect of specific membrane conductance $G_{pas}$ on voltage decay when the product of $R_a$ and $G_{pas}$ is fixed: across 19 Y3 neurons simulated, the voltage decay measured as a ratio of the voltage in the medulla region and the lobula region is the same across 0.09, 0.13 and 0.2 mS/cm$^2$. For (**c-d**), $\rho = 1/(4 \times R_a \times G_{pas})$ is fixed to 12.5 cm, and describes how $R_a$ and $G_{pas}$ affects voltage attenuation according to cable theory. The space constant of passive cable theory is $\lambda = \sqrt{\rho d}$ where d is the diameter of the cable.

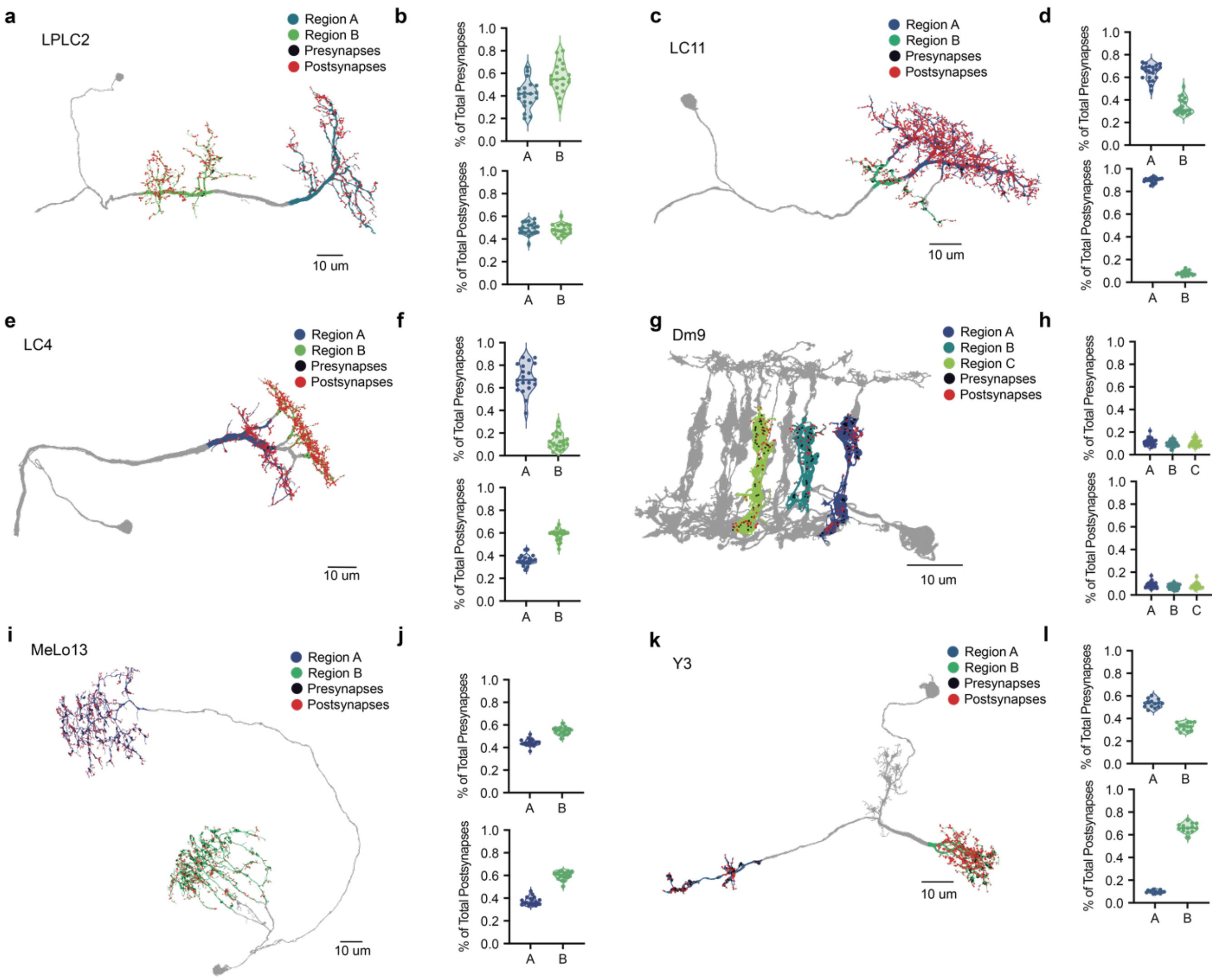


**Extended Data Figure 8. Distribution of postsynapses and presynapses across regions. a**, Representative nanoscale reconstruction of an LPLC2 neuron, with regions highlighted as marked. Presynapses are marked with red dots, and postsynapses are marked with black dots. **b,** Quantification of presynapses (top) and postsynapses (bottom) in regions A and B across reconstructed LPLC2 neurons (*n* = 20) as a percentage of total presynapses and postsynapses per cell. Data shown as violin plots and reported as median [25$^{th}$ percentile, 75$^{th}$ percentile]. Top: Region A, 42.0% [34.3%, 48.8%]; Region B, 54.9% [48.1%, 65.3%]. Bottom: Region A, 49.5% [45.8%, 53.6%]; Region B 48.0% [44.6%, 52.3%]. **c-d**, as in **a-b** for LC11 (*n* = 20). **d**, Top: Region A, 66.5% [59.9%, 69.8%]; Region B, 31.2% [26.5%, 39.9%]. Bottom: Region A, 90.8% [89.1%, 91.5%]; Region B 7.7% [6.9%, 8.9%]. **e-f**, as in **a-b** for LC4 (*n* = 20). **f**, Top: Region A, 67.0% [60.7%, 80.7%]; Region B, 11.4% [7.9%, 18.5%]. Bottom: Region A, 35.1% [33.4%, 37.8%]; Region B, 59.9% [54.7%, 60.7%]. **g-h**, as in **a-b**, for Dm9 (*n* = 19). **h**, Top: Region A, 11.1% [9.2%, 12.8%]; Region B, 10.3% [8.0%, 12.1%]; Region C, 9.7% [8.3%, 12.4%]. Bottom: Region A 8.9% [7.6%, 10.5%]; Region B, 7.3% [5.6%, 9.7%]; Region C, 7.6% [6.2%, 9.4%]. **i-j**, as in **a-b** for MeLo13 (*n* = 20). **j**, Top: Region A, 43.8% [42.7%, 46.4%]; Region B, 55.2% [52.0%, 56.6%]. Bottom: Region A, 36.1% [34.5%, 39.5%]; Region B, 60.8% [58.3%, 62.8%]. **k-l**, as in **a-b**, for Y3 (*n* = 13). **l**, Top: Region A, 53.4% [50.6%, 56.3%]; Region B, 33.0% [29.7%, 36.5%]. Bottom: Region A, 10.0% [9.1%, 11.0%]; Region B, 66% [63.2%, 69.4%].

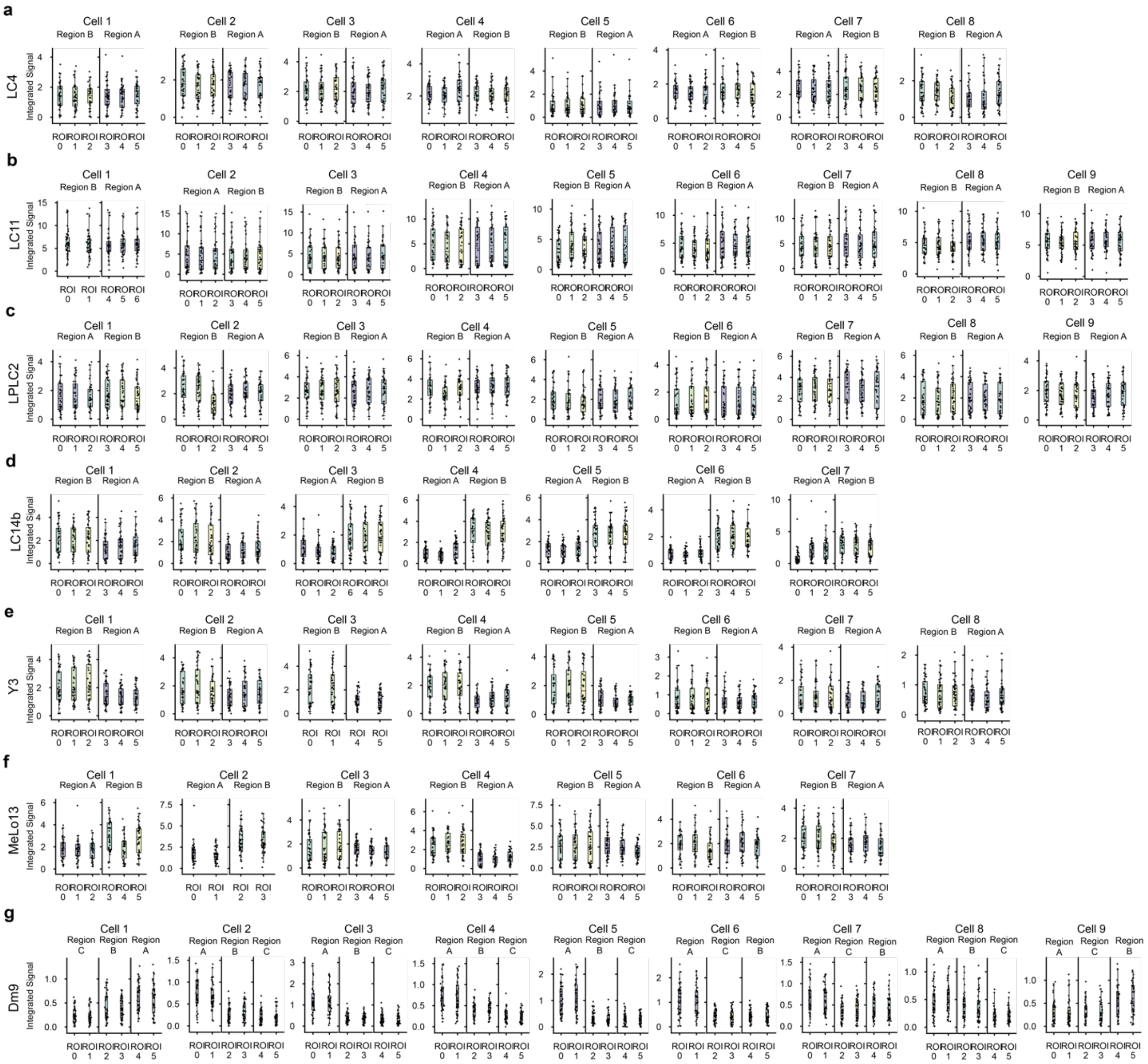

**Extended Data Figure 9. Voltage distributions within regions. a,** Integrated signal across individual ROIs in Region A and Region B for all imaged LC4 neurons. For each ROI, the integrated signal for the –ΔF/F trace was computed for every trial. Single dots represent individual trials; boxplots show the median and the interquartile range (25th-75th percentiles). **b**, Same as (**a**) for LC11. **c**, Same as (**a**) for LPLC2. **d**, Same as (**a**) for LC14b. **e**, Same as (**a**) for Y3. **f**, Same as (**a**) for MeLo13. **g**, Same as (**a**) for Dm9 (including three regions).

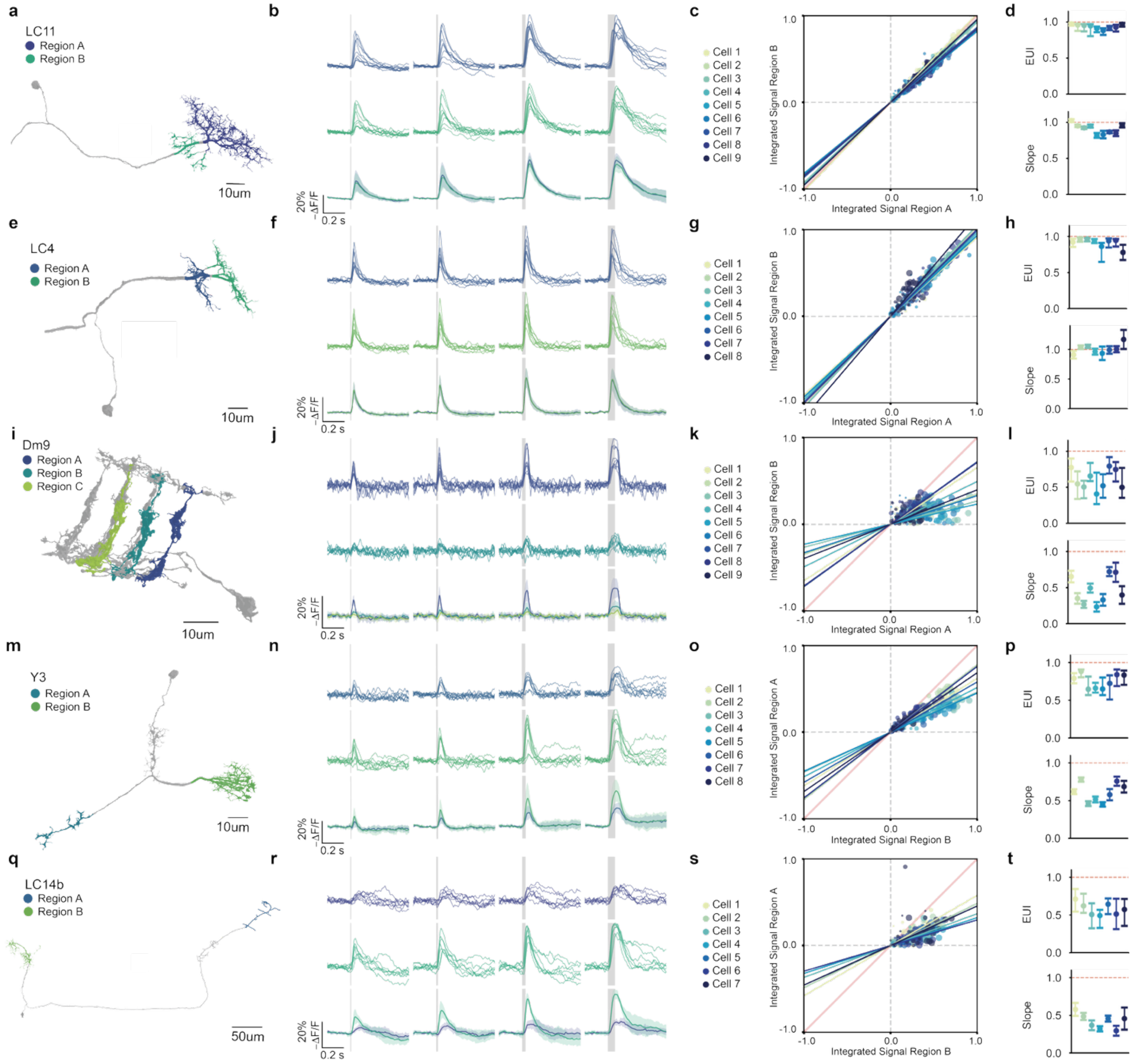

**Extended Data Figure 10. Visual neurons span a continuum of electrical properties. a,** Nanoscale reconstruction of an LC11 neuron with Regions A and B highlighted. **b**, Stimulus evoked responses of measured LC11 neurons (*n* = 9 cells, 7 flies) across stimulus durations. Stimuli are shown from left to right in order of increasing duration (8, 16, 32, 64 ms), as indicated by grey shading. Top: each line represents the average –ΔF/F response in Region A for a recorded LC11 neuron over 10 stimulus presentations. Middle: Corresponding average –ΔF/F responses in Region B. Bottom: overlay of mean responses in Region A and Region B across all recorded LC11 neurons. Colored shading denotes standard deviation in Region A and Region B. **c**, Integrated signal in Region B plotted against Region A overall measured LC11 neurons and trials. Each color represents trials from an individual neuron. Data are plotted as described in (**Fig. 5d)**, with zero-intercept linear regression fits overlaid for each cell. **d**, Quantification of EUI and slope of zero-intercept linear regressions across measured LPLC2 neurons. Top: quantification of EUI for each LC11 neuron over all trials. Data shown as median with 25th -75th percentile error bars. Population median = 0.95. Bottom: quantification of zero-intercept linear regressions for each LC11 neuron over all trials. Data shown as slope with the 95% confidence interval in error bars. Population median = 0.93. **e-h**, same as (**a-d**) for LC4 (*n* = 8 cells, 7 flies). **h**, Top: population median = 0.94. Bottom: population median = 1.0. **i-l**, same as (**a-d**) for Dm9 Regions A and B. **l**, Top: population median = 0.55. Bottom: population median = 0.40. **m-p**, same as (**a-d**) for Y3. **p**, Top: population median = 0.76. Bottom: population median = 0.60. **q-t**, Same as (**a-d**) for LC14b. **t**, Top: population median = 0.57. Bottom: population median = 0.46. For quantification of individual cells across all cell types see **Extended Data Table.**

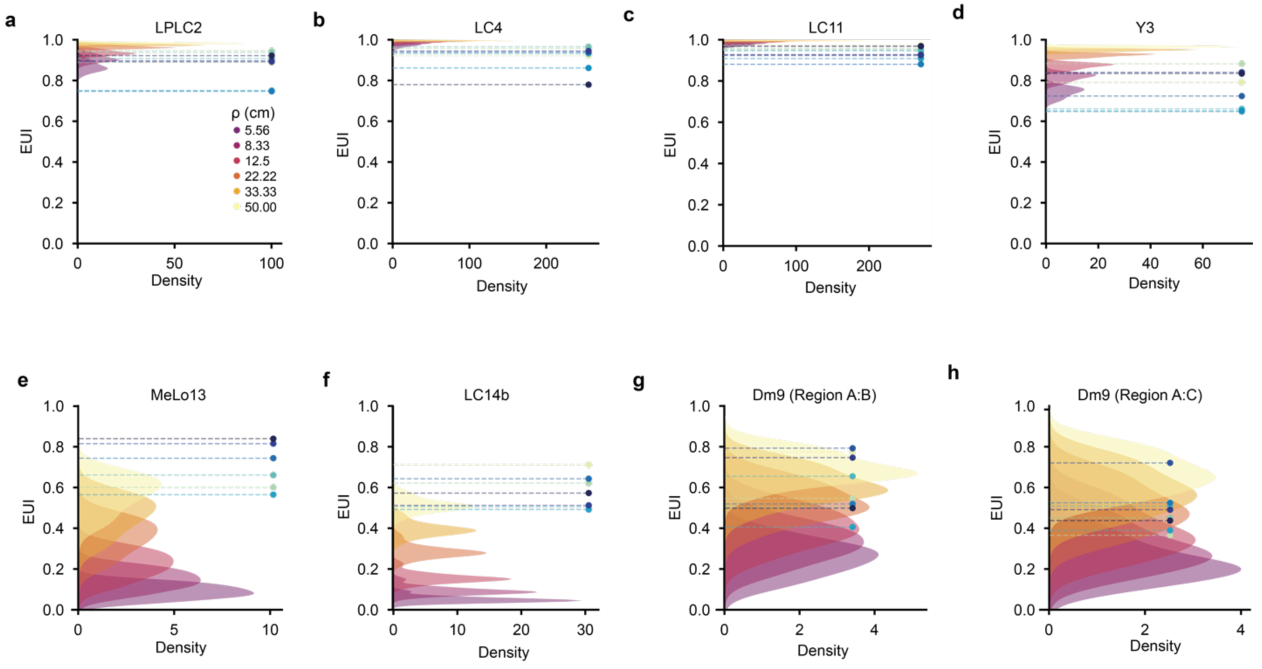


**Extended Data Figure 11. Comparison of voltage distributions between modeled and measured neurons. a,** Distribution of modeling results from passive simulation of reconstructed LPLC2 neurons (*n* = 20). Single neuronal reconstructions were simulated in NEURON across a range of biophysical parameters. To calculate the EUI for each simulation and each reconstruction, the average voltage recorded across Region A and B was used. The distribution of EUI values across biophysical parameters is displayed as a continuous probability density function. Distributions corresponding to different biophysical parameter sets are color-coded as indicated. Experimentally measured median EUIs for each LPLC2 cell (*n* = 9) are overlaid as dashed lines (see **Fig. 5** and **Extended Data Figure 10**). **b**, Same as (**a**) for LC4 (modeled, *n* = 20; measured, *n* = 8). **c**, Same as (**a**) for LC11 (modeled, *n* = 20; measured, *n* = 9). **d**, Same as (**a**) for Y3 (modeled, *n* = 13; measured *n* = 8). **e**, Same as (**a**) for MeLo13 (modeled, *n* = 20; measured, *n* = 7). **f**, Same as (**a**) for LC14b (modeled, *n* = 18; measured, *n* = 7). **g**, Same as (**a**) for Regions A and B of Dm9 (modeled, *n* = 19; measured, *n* = 9). **h**, Same as (**a**) for Regions A and C of Dm9 (modeled, *n* = 19; measured, *n* = 9).

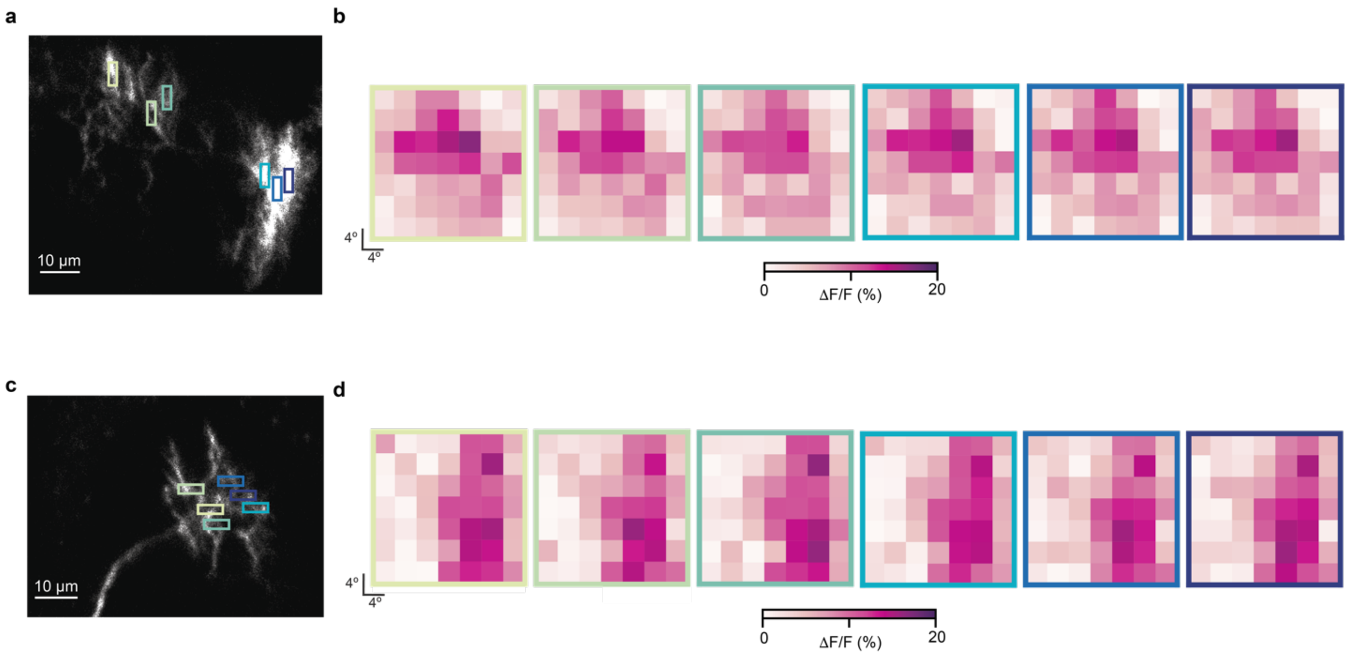


**Extended Data Figure 12. LPLC2 and LC4 display uniform spatial receptive fields across neurites. a**, Imaged LPLC2 neuron with overlaid ROIs. **b,** Spatial tuning maps for each ROI, measured using fine receptive field mapping (see **Fig. 3**). **c-d**, Same as **(a-b)** for LC4.

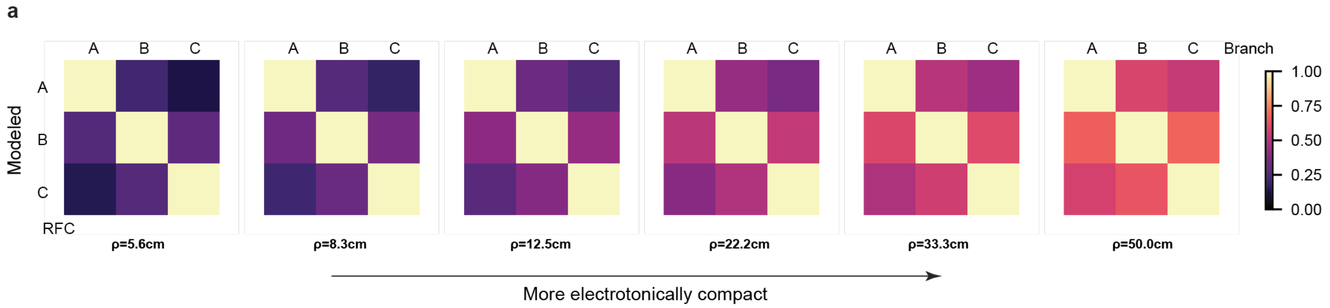


**Extended Data Figure 13. Simulated confusion matrices of Dm9 across biophysical parameters.** Simulated confusion matrices of Dm9 neuron type across relevant biophysical parameters. Pixel (i, j) is the integrated voltage signal of branch i to the stimulation of branch j normalized by diagonal pixel (i, i).

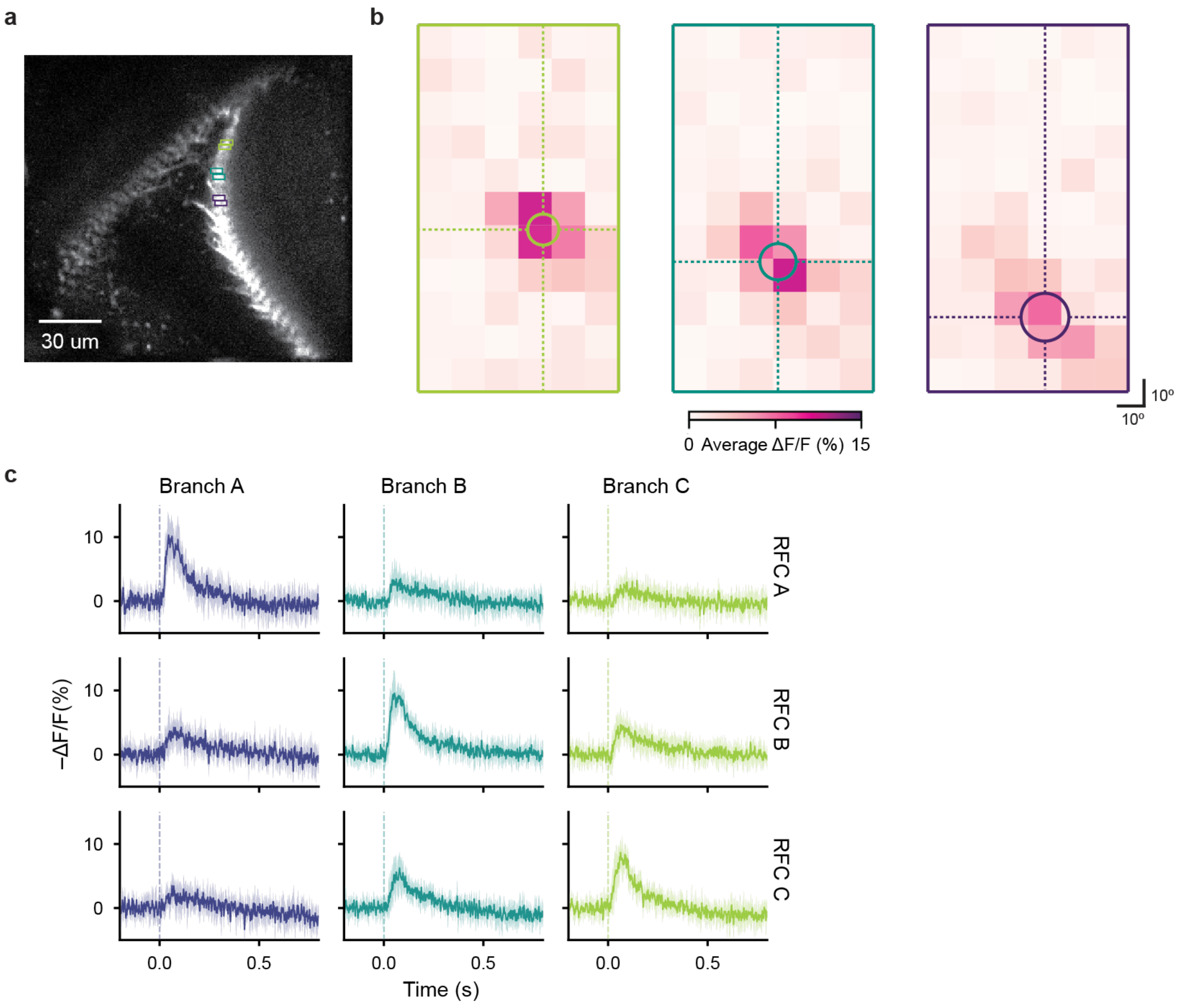


**Extended Data Figure 14. Voltage compartmentalization in CT1 produces branch-specific spatial-tuning of receptive field. a**, An imaged CT1 neuron with overlaid ROI groups. **b**, Spatial tuning maps for each ROI group in (**a**). A 2D Gaussian was fit to –ΔF/F responses to extract the center and extent of the peak response, visualized as overlaid ellipses. **c**, Branch-specific CT1 responses across all measured neurons ($n$ = 7 cells, 6 flies). The receptive field center of each CT1 branch was mapped in a 60º × 110º grid, and subsequently a dark spot stimulus (r = 10º) was presented at the receptive field center of each ROI group. Responses were recorded across all branches, and the average response per branch, per neuron was calculated. Data are shown as the average –ΔF/F response per branch across CT1 neurons; colored shading denotes standard error of the mean across cells.

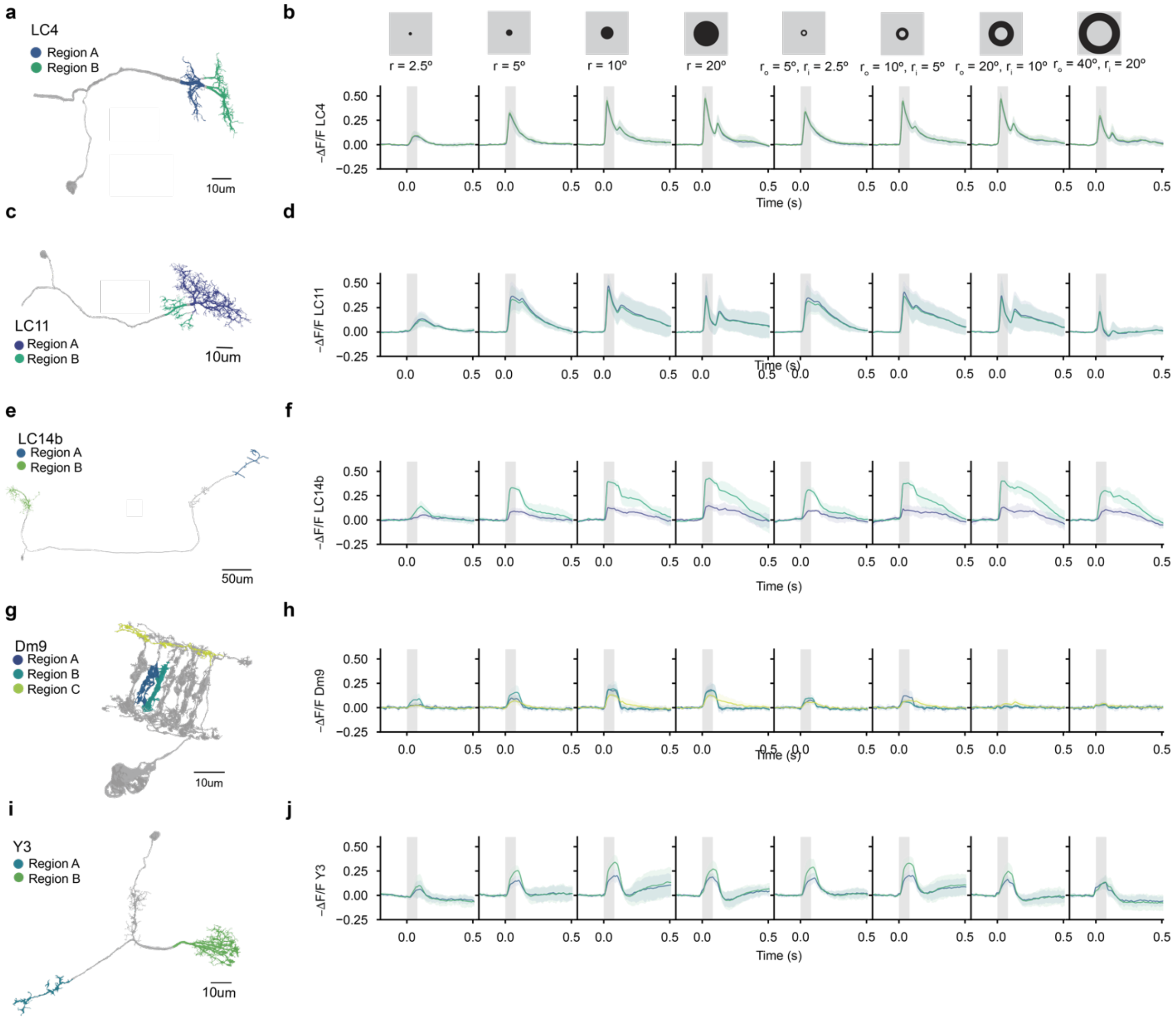


**Extended Data Figure 15. Responses across visual neurons to additional spots and annuli. a**, Nanoscale reconstruction of an LC4 neuron with Regions A and B highlighted (as in Extended Data Figure 9). **b**, Responses across LC4 neurons (*n* = 8 cells, 7 flies) for Regions A and B to dark spots and annuli of increasing radii. Each stimulus was presented over 10 trials, and the average response per region, per neuron was calculated. Data shown as average –ΔF/F response per region across measured LC4 neurons; region responses are color-coded as in (**a**); colored shading represents standard error of the mean (S.E.M.); grey shading denotes stimulus duration. **c-d**, Same as (**a-b**) for LC11 (*n* = 9 cells, 7 flies). **e-f**, Same as (**a-b**) for LC14b (*n* = 7 cells, 7 flies). **g-h**, Same as (**a-b**) for Dm9 (*n* = 9 cells, 9 flies). **i-j**, Same as (**a-b**) for Y3 (*n* = 7 cells, 5 flies).

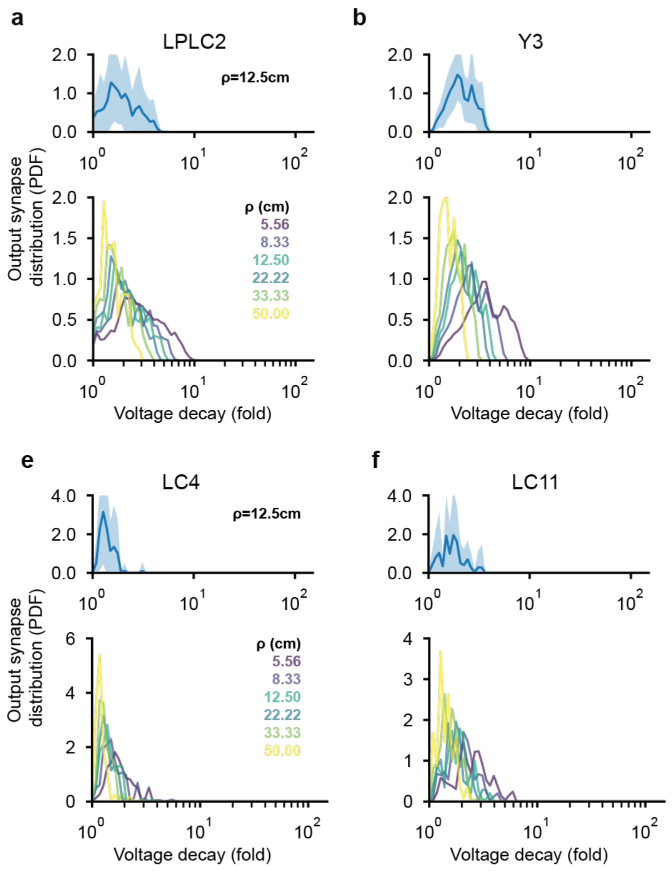


**Extended Data Figure 16. Distribution of output synapses over voltage decay across cell types. a**, Top: the probability density function (PDF) of the distribution of output synapses across voltage decay of simulated LPLC2 neurons. *n*=20 neurons simulated with the space constant factor ρ=12.5 cm. Shading denotes the standard deviation. Bottom: average PDF of the distribution of output synapses across voltage decay of simulated LPLC2 neurons at different ρ values. **b**, same as (**a**), for simulated Y3 neurons (*n*=18). **c**, same as (**a**), for simulated MeLo13 neurons (*n*=20). **d**, same as (**a**), for simulated Dm9 neurons (*n*=20). **e**, same as (**a**), for simulated LC4 neurons (*n*=20). **f**, same as (**a**), for simulated LC11 neurons (*n*=20).

| LPLC2 | EUI median [25th -75th %] | Slope 95% CI [lower bound, upper bound] |
|---|---|---|
| Cell 1 | 0.888 [0.827, 0.952] | 1.052 [0.988 - 1.117], Adjusted $R^2$= 0.964, ****$P < 0.0001$ |
| Cell 2 | 0.947 [0.908, 0.978] | 0.950 [0.901 - 1.000], Adjusted $R^2$= 0.974, ****$P < 0.0001$ |
| Cell 3 | 0.939 [0.899, 0.983] | 0.982 [0.938 - 1.025], Adjusted $R^2$= 0.981, ****$P < 0.0001$ |
| Cell 4 | 0.908 [0.805, 0.966] | 0.905 [0.839 - 0.971], Adjusted $R^2$= 0.951, ****$P < 0.0001$ |
| Cell 5 | 0.750 [0.636, 0.869] | 0.923 [0.805 - 1.042], Adjusted $R^2$= 0.860, ****$P < 0.0001$ |
| Cell 6 | 0.747 [0.444, 0.914] | 0.945 [0.798 - 1.092], Adjusted $R^2$= 0.807, ****$P < 0.0001$ |
| Cell 7 | 0.892 [0.845, 0.954] | 0.941 [0.872 - 1.009], Adjusted $R^2$= 0.951, ****$P < 0.0001$ |
| Cell 8 | 0.896 [0.666, 0.964] | 1.010 [0.948 - 1.071], Adjusted $R^2$= 0.965, ****$P < 0.0001$ |
| Cell 9 | 0.922 [0.891, 0.961] | 1.020 [0.971 - 1.069], Adjusted $R^2$= 0.978, ****$P < 0.0001$ |
| **Dm9 (A-C)** | **EUI median [25th -75th %]** | **Slope 95% CI [lower bound, upper bound]** |
| Cell 1 | 0.525 [0.350, 0.638] | 0.304 [0.239 - 0.370], Adjusted $R^2$= 0.688, ****$P < 0.0001$ |
| Cell 2 | 0.362 [0.172, 0.562 | 0.234 [0.168 - 0.299], Adjusted $R^2$= 0.562, ****$P < 0.0001$ |
| Cell 3 | 0.447 [0.231, 0.692] | 0.209 [0.135 - 0.284], Adjusted $R^2$= 0.438, ****$P < 0.0001$ |
| Cell 4 | 0.504 [0.288, 0.608] | 0.279 [0.224 - 0.335], Adjusted $R^2$= 0.717, ****$P < 0.0001$ |
| Cell 5 | 0.386 [0.211, 0.703] | 0.211 [0.145 - 0.278], Adjusted $R^2$= 0.502, ****$P < 0.0001$ |
| Cell 6 | 0.521 [0.302, 0.745] | 0.323 [0.250 - 0.397], Adjusted $R^2$= 0.661, ****$P < 0.0001$ |
| Cell 7 | 0.717 [0.555, 0.892] | 0.540 [0.449 - 0.632], Adjusted $R^2$= 0.781, ****$P < 0.0001$ |
| Cell 8 | 0.487 [0.358, 0.702 | 0.319 [0.228 - 0.410], Adjusted $R^2$= 0.550, ****$P < 0.0001$ |
| Cell 9 | 0.434 [0.220, 0.780] | 0.355 [0.208 - 0.502], Adjusted $R^2$= 0.364, ****$P < 0.0001$ |
| **MeLo13** | **EUI median [25th -75th %]** | **Slope 95% CI [lower bound, upper bound]** |
| Cell 1 | 0.833 [0.642, 0.893] | 0.637 [0.555 - 0.718], Adjusted $R^2$= 0.865, ****$P < 0.0001$ |
| Cell 2 | 0.596 [0.493, 0.795] | 0.479 [0.410 - 0.549], Adjusted $R^2$= 0.829, ****$P < 0.0001$ |
| Cell 3 | 0.656 [0.427, 0.852] | 0.568 [0.416 - 0.721], Adjusted $R^2$= 0.582, ****$P < 0.0001$ |
| Cell 4 | 0.560 [0.313, 0.675] | 0.355 [0.295 - 0.414], Adjusted $R^2$= 0.782, ****$P < 0.0001$ |
| Cell 5 | 0.739 [0.629, 0.842] | 0.675 [0.562 - 0.788], Adjusted $R^2$ = 0.783, ****$P < 0.0001$ |
| Cell 6 | 0.810 [0.721, 0.892] | 0.888 [0.745 - 1.031], Adjusted $R^2$= 0.796, ****$P < 0.0001$ |

| | | |
|---|---|---|
| Cell 7 | 0.834 [0.785, 0.895] | 0.763 [0.679 - 0.848], Adjusted $R^2$= 0.893, ****$P < 0.0001$ |
| **LC11** | **EUI median [25th -75th %]** | **Slope 95% CI [lower bound, upper bound]** |
| Cell 1 | 0.974 [0.949, 0.988] | 1.024 [1.004 – 1.044], Adjusted $R^2$= 0.996, ****$P < 0.0001$ |
| Cell 2 | 0.959 [0.877, 0.973] | 0.952 [0.935 - 0.968], Adjusted $R^2$= 0.997, ****$P < 0.0001$ |
| Cell 3 | 0.953 [0.870, 0.971] | 0.925 [0.907 - 0.943], Adjusted $R^2$= 0.996, ****$P < 0.0001$ |
| Cell 4 | 0.946 [0.804, 0.971] | 0.948 [0.927 - 0.970], Adjusted $R^2$= 0.995, ****$P < 0.0001$ |
| Cell 5 | 0.908 [0.858, 0.936] | 0.820 [0.787 - 0.853], Adjusted $R^2$= 0.984, ****$P < 0.0001$ |
| Cell 6 | 0.881 [0.821, 0.914] | 0.831 [0.780 - 0.883], Adjusted $R^2$= 0.964, ****$P < 0.0001$ |
| Cell 7 | 0.924 [0.887, 0.946] | 0.867 [0.849 - 0.885], Adjusted $R^2$= 0.996, ****$P < 0.0001$ |
| Cell 8 | 0.927 [0.868, 0.962] | 0.850 [0.810 - 0.890], Adjusted $R^2$= 0.979, ****$P < 0.0001$ |
| Cell 9 | 0.969 [0.936, 0.988] | 0.959 [0.930 - 0.989], Adjusted $R^2$= 0.991, ****$P < 0.0001$ |
| **LC4** | **EUI median [25th -75th %]** | **Slope 95% CI [lower bound, upper bound]** |
| Cell 1 | 0.922 [0.855, 0.969] | 0.910 [0.845 - 0.974], Adjusted $R^2$= 0.953, ****$P < 0.0001$ |
| Cell 2 | 0.957 [0.926, 0.987] | 1.037 [1.007 - 1.068], Adjusted $R^2$= 0.992, ****$P < 0.0001$ |
| Cell 3 | 0.965 [0.931, 0.978] | 1.051 [1.024 - 1.079], Adjusted $R^2$= 0.994, ****$P < 0.0001$ |
| Cell 4 | 0.932 [0.908, 0.965] | 0.963 [0.912 - 1.013], Adjusted $R^2$= 0.974, ****$P < 0.0001$ |
| Cell 5 | 0.861 [0.648, 0.911] | 0.934 [0.821 - 1.048], Adjusted $R^2$= 0.876, ****$P < 0.0001$ |
| Cell 6 | 0.939 [0.846, 0.973] | 0.990 [0.927 - 1.054], Adjusted $R^2$= 0.962, ****$P < 0.0001$ |
| Cell 7 | 0.945 [0.861, 0.965] | 1.003 [0.952 - 1.055], Adjusted $R^2$= 0.974, ****$P < 0.0001$ |
| Cell 8 | 0.779 [0.674, 0.884] | 1.169 [1.008 - 1.329], Adjusted $R^2$= 0.843, ****$P < 0.0001$ |
| **Dm9 (A-B)** | **EUI median [25th -75th %]** | **Slope 95% CI [lower bound, upper bound]** |
| Cell 1 | 0.774 [0.578, 0.898] | 0.653 [0.572 - 0.734], Adjusted $R^2$ = 0.870,****$P < 0.0001$ |
| Cell 2 | 0.547 [0.337, 0.720] | 0.349 [0.276 - 0.423], Adjusted $R^2$ = 0.696, ****$P < 0.0001$ |
| Cell 3 | 0.504 [0.347, 0.590] | 0.267 [0.218 - 0.316], Adjusted $R^2$ = 0.750, ****$P < 0.0001$ |
| Cell 4 | 0.655 [0.585, 0.837] | 0.493 [0.431 - 0.555], Adjusted $R^2$ = 0.864, ****$P < 0.0001$ |
| Cell 5 | 0.406 [0.266, 0.702] | 0.233 [0.167 - 0.298], Adjusted $R^2$ = 0.556, ****$P < 0.0001$ |
| Cell 6 | 0.520 [0.354, 0.680] | 0.330 [0.249 - 0.411], Adjusted $R^2$ = 0.625, ****$P < 0.0001$ |

| | | |
|---|---|---|
| Cell 7 | 0.793 [0.690, 0.917] | 0.721 [0.658 - 0.784], Adjusted $R^2$ = 0.931, ****$P < 0.0001$ |
| Cell 8 | 0.746 [0.579, 0.848] | 0.713 [0.579 - 0.847], Adjusted $R^2$ = 0.742, ****$P < 0.0001$ |
| Cell 9 | 0.499 [0.353, 0.768] | 0.397 [0.274 - 0.519], Adjusted $R^2$ = 0.511, ****$P < 0.0001$ |
| **Y3** | **EUI median [25th -75th %]** | **Slope 95% CI [lower bound, upper bound]** |
| Cell 1 | 0.791 [0.726, 0.860] | 0.619 [0.586 - 0.652], Adjusted $R^2$= 0.973, ****$P < 0.0001$ |
| Cell 2 | 0.882 [0.805, 0.908] | 0.779 [0.751 - 0.806], Adjusted $R^2$= 0.988, ****$P < 0.0001$ |
| Cell 3 | 0.647 [0.569, 0.814] | 0.461 [0.430 - 0.492], Adjusted $R^2$= 0.956, ****$P < 0.0001$ |
| Cell 4 | 0.660 [0.602, 0.732] | 0.515 [0.477 - 0.553], Adjusted $R^2$= 0.949, ****$P < 0.0001$ |
| Cell 5 | 0.650 [0.569, 0.802] | 0.451 [0.421 - 0.480], Adjusted R2= 0.959, ****P < 0.0001 |
| Cell 6 | 0.723 [0.510, 0.831] | 0.579 [0.504 - 0.654], Adjusted $R^2$= 0.859, ****$P < 0.0001$ |
| Cell 7 | 0.841 [0.689, 0.908] | 0.758 [0.698 - 0.818], Adjusted $R^2$= 0.942, ****$P < 0.0001$ |
| Cell 8 | 0.834 [0.704, 0.894] | 0.686 [0.608 - 0.763], Adjusted $R^2$= 0.888, ****$P < 0.0001$ |
| **LC14b** | **EUI median [25th -75th %]** | **Slope 95% CI [lower bound, upper bound]** |
| Cell 1 | 0.711 [0.545, 0.845] | 0.579 [0.493 - 0.665], Adjusted $R^2$= 0.821, ****$P < 0.0001$ |
| Cell 2 | 0.621 [0.528, 0.783] | 0.486 [0.438 - 0.533], Adjusted $R^2$= 0.915, ****$P < 0.0001$ |
| Cell 3 | 0.506 [0.324, 0.655 | 0.368 [0.303 - 0.432], Adjusted $R^2$= 0.766, ****$P < 0.0001$ |
| Cell 4 | 0.492 [0.329, 0.568] | 0.322 [0.287 - 0.356], Adjusted $R^2$= 0.898, ****$P < 0.0001$ |
| Cell 5 | 0.643 [0.512, 0.720] | 0.456 [0.417 - 0.496], Adjusted $R^2$= 0.932, ****$P < 0.0001$ |
| Cell 6 | 0.512 [0.312, 0.716] | 0.296 [0.230 - 0.361], Adjusted $R^2$= 0.672, ****$P < 0.0001$ |
| Cell 7 | 0.573 [0.355, 0.714] | 0.457 [0.310 - 0.604], Adjusted $R^2$= 0.491, ****$P < 0.0001$ |

**Extended Data Table 1. Summary of EUI and linear regressions for individual Cells.** EUI calculated over all individual trials for each cell (n=40) and reported as median with 95% confidence intervals. Constrained ordinary least squares linear regression performed over all trials for each cell (n=40, degree of freedom = 39). Reported as slope with 95% confidence intervals.
